# Kinetic Pathways of Topology Simplification by Type-II Topoisomerases in Knotted Supercoiled DNA

**DOI:** 10.1101/383901

**Authors:** Riccardo Ziraldo, Andreas Hanke, Stephen D. Levene

**Author notes:** To whom correspondence should be addressed. Correspondence may also be addressed to Andreas Hanke. These authors contributed equally to the work as first authors (Riccardo Ziraldo, Andreas Hanke).

## Abstract

The topological state of covalently closed, double-stranded DNA is defined by the knot type *K* and the linking-number difference Δ*Lk* relative to unknotted relaxed DNA. DNA topoisomerases are essential enzymes that control the topology of DNA in all cells. In particular, type-II topoisomerases change both *K* and Δ*Lk* by a duplex-strand-passage mechanism and have been shown to simplify the topology of DNA to levels below thermal equilibrium at the expense of ATP hydrolysis. It remains a puzzle how small enzymes are able to preferentially select strand passages that result in topology simplification in much larger DNA molecules. Using numerical simulations, we consider the non-equilibrium dynamics of transitions between topological states (*K*, Δ*Lk*) in DNA induced by type-II topoisomerases. For a biological process that delivers DNA molecules in a given topological state (*K*,Δ*Lk*) at a constant rate we fully characterize the pathways of topology simplification by type-II topoisomerases in terms of stationary probability distributions and probability currents on the network of topological states (*K*,Δ*Lk*). In particular, we observe that type-II topoisomerase activity is significantly enhanced in DNA molecules that maintain a supercoiled state with constant torsional tension. This is relevant for bacterial cells in which torsional tension is maintained by enzyme-dependent homeostatic mechanisms such as DNA-gyrase activity.

## INTRODUCTION

The topological state of covalently closed, double-stranded DNA is defined by the knot type, *K*, and the linking number, *Lk*. DNA topoisomerases play a critical role in controlling the topology of double-stranded DNA through torsional relaxation and supercoiling, decatenation of interlocked DNA duplexes, and elimination of knotted DNA-recombination products, which cannot support transcription and replication (1–5). Supercoiling is quantitatively defined in terms of the linking-number difference relative to relaxed DNA, Δ*Lk* = *Lk* − *Lk*_0_, rather than *Lk* itself; here, *Lk*_0_ = *N*/*h*_0_ where *N* is the number of DNA base pairs in the DNA molecule and *h*_0_ is the number of base pairs per helical turn in topologically relaxed DNA.

DNA topoisomerases are divided into two classes, type-I and type-II, corresponding to mechanisms that involve cleavage of one or both DNA strands, respectively (6). Type-I enzymes regulate the torsional tension in double-stranded DNA by changing Δ*Lk* exclusively whereas type-II enzymes can change both *K* and Δ*Lk* by passing one duplex DNA segment through another. Torsional relaxation of DNA is energetically favorable and can be performed by ATP-independent enzymes, such as topoisomerase I, and by DNA gyrase in the absence of ATP (3). All topoisomerases can remove supercoils from DNA, but DNA gyrase can also introduce negative supercoils into DNA at the expense of ATP hydrolysis. All type-II enzymes require ATP hydrolysis to perform duplex-segment passage, in particular type-IIA enzymes such as bacterial topoisomerase IV and eukaryotic topoisomerase II (7). This cofactor requirement was poorly understood until Rybenkov *et al*. showed in 1997 that type-II topoisomerases selectively perform strand passages that reduce the steady-state fraction of knotted or catenated, torsionally relaxed plasmid DNAs to levels 80 times below that at thermal equilibrium (8). In particular, the width of the Δ*Lk* distribution for torsionally relaxed plasmid DNAs acted on by type-IIA topoisomerases was found to be narrower, i.e., less supercoiled, than that observed with ATP-independent enzymes (8). Thus, type-II topoisomerases use the free energy of ATP hydrolysis to drive the system away from thermal equilibrium. However, the puzzle remains how a relatively small enzyme is able to preferentially select strand passages that lead to unknotting rather than to formation of knots in large DNA molecules because the topological state of DNA is a property of the entire molecule that cannot be determined by local DNA-enzyme interactions.

Since the seminal work by Rybenkov *et al*. several models have been suggested to explain how ATP-hydrolysis-driven type-II topoisomerases can selectively lower the frequency of DNA knotting (8–14). These models are generally based on geometric or kinetic mechanisms that increase the probability of strand-passage reactions and result in topology simplification from an initial state. The most successful model that has emerged from these studies is the model of a hairpin-like gate (G) segment, where the type-II enzyme strongly bends the G-segment DNA and accepts for passage only a transfer (T) segment from the inside to the outside of the hairpin-formed G segment (Figure 1A) (9, 15). For torsionally unconstrained (nicked) DNA, the model predicts a large decrease in the steady-state proportion of knots and catenanes relative to those at equilibrium, although it is insufficient to explain the magnitude of the effect observed with torsionally relaxed DNA plasmids (9, 15). Indeed, strong (~150°) protein-induced bending of the G segment, as required by the model, is observed in a co-crystal structure of yeast topoisomerase II with G-segment DNA (Figure 1B) (16). Experimental AFM measurements are consistent with bend angles between 94° and 100°, whereas FRET measurements suggest somewhat larger bend angles of 126° and 140° (17).

**Figure 1.**
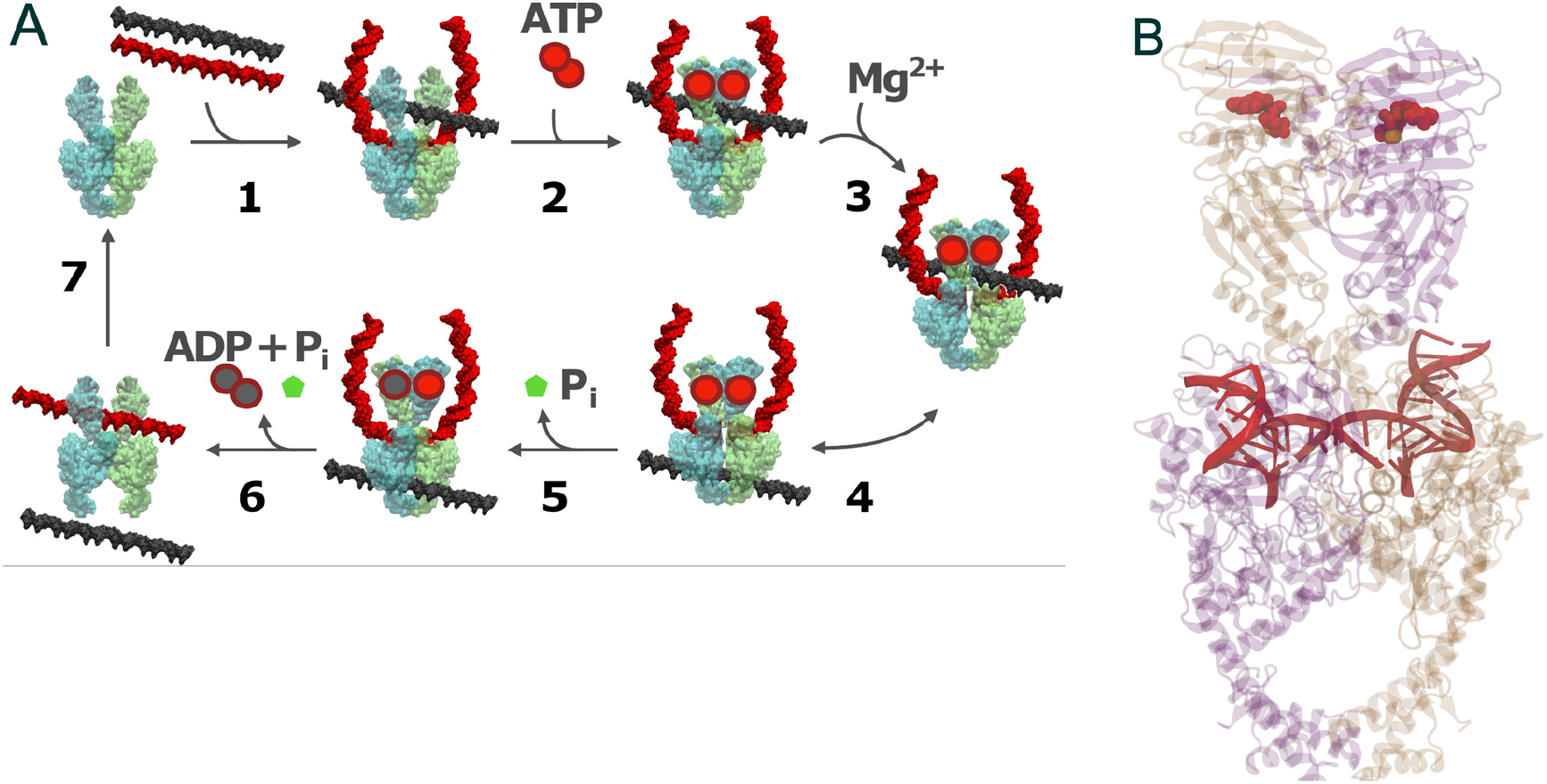
(**A**) Mechanistic details of duplex-DNA passage in type-II topoisomerases. Step 1: enzyme binding to gate (G) segment in duplex DNA, followed by an encounter with transfer (T) segment; step 2: binding of two ATP molecules seals the gate; step 3: cleavage of the G segment duplex, catalyzed by the binding of 2 Mg^2+^ ions; step 4: passage of the T segment through the G segment; step 5: hydrolysis of the first ATP molecule releases a phosphate group and reseals the G segment strand; step 6: hydrolysis of the second ATP molecule dismantles the complex releasing both DNA strands; step 7: the enzyme resets to its original conformation. (**B**) Structure of yeast type-II topoisomerase dimer bound to a doubly nicked 34-mer duplex DNA (PDB 4GFH) and AMP-PNP. The DNA is bent 160° via interactions with an invariant isoleucine (49).

Another group of studies did not directly address mechanisms of topoisomerase action but considered the probability distribution *P*(*K*, Δ*Lk*), and distributions derived therefrom, at thermal equilibrium (10, 18–20). This distribution is directly related to the free-energy landscape *F*(*K*, Δ*Lk*) = −*k_B_T* ln*P*(*K*, Δ*Lk*) where *T* is the temperature and *k_B_* is Boltzmann’s constant. The distribution *P*(*K*,Δ*Lk*) corresponds to a phantom-chain ensemble where the DNA molecules are free to explore all topological states (*K*,Δ*Lk*) at thermal equilibrium, referred to here as the equilibrium segment-passage (ESP) ensemble (21). Characterization of *P*(*K*,Δ*Lk*) therefore yields important insight about the most likely relaxation path of a given DNA knot by a hypothetical topoisomerase that lacks any bias towards topology simplification and is driven only by the topological free-energy gradient. Indeed, the actual extent of bias for an ATP-driven type-II enzyme in favor of unknotting can only be quantified if we know the probability of acting in the absence of any bias, corresponding to topoisomerase action in absence of ATP hydrolysis. The system’s behavior at thermal equilibrium thus provides a necessary reference state for investigating mechanisms of topoisomerase activity such as chirality bias (22–24).

Motivated by the fact that type-II enzymes drive the system away from equilibrium, we investigate a model of topoisomerase activity based on a network of topological states (*K*,Δ*Lk*) of circular DNAs with knot type *K* and linking number difference Δ*Lk* in which the dynamics of transitions between states (*K*,Δ*Lk*) mediated by type-II enzymes is described by a chemical master equation. Previous studies showed the existence of unknotting/unlinking pathways followed by type-II topoisomerases that stepwise progressively reduce the topological complexity of knotted/catenated molecules (25, 26). The main goal of our study is to identify significant pathways along which topology simplification by type-II enzymes occurs in terms of non-equilibrium steady states (NESSs) for the network (*K*,Δ*Lk*). We also quantify type-II topoisomerase activity for a hairpin-like G segment compared to a straight (unbent) G segment. To address these questions we generated a large set of equilibrium ensembles of knotted and supercoiled 6-kbp DNAs by Monte Carlo simulations to find transition rates and NESS parameters in the network of topological states (*K*,Δ*Lk*). Our analytical approach can be thought of as a two-level model. At the top (macroscopic) level, the model uses topological states as the variable, so it allows DNA-topology transitions between states (*K*,Δ*Lk*) according to a chemical master equation. This state space is therefore an integer lattice and the transitions between states occur with rates that are computed from an explicit, coarse-grained polymer model, which accounts for microscopic states. The master-equation formulation allows one to compute the occupancies of the different macrostates, including their dynamics. In principle, other mesoscopic models for knotted supercoiled DNA can be used to capture the underlying microscopic behavior of the system, such as those obtained from Brownian or molecular dynamics (27).

A novel feature of our model is the capability to dynamically account for processes that generate complex knots extraneously, either *in vitro* or *in vivo*. The favorable unknotting pathways were determined in terms of universal NESS probabilities and probability currents, derived from transition rates. The idea of an induced probability current stems from the presence of an idealized source of complex knots. For example, type-II enzymes crucially maintain the integrity of genomic DNA during transcription and replication, requiring relaxation of (+) supercoils that build up ahead of RNA and DNA polymerases (28, 29). High local concentrations of type-II enzyme molecules near the boundary of a transcription bubble or ahead of a replication fork could therefore increase the probability of knotting through stochastic duplex-segment passage. *In vitro*, type-IIA enzymes efficiently generate not-trivial knots through processes that facilitate intramolecular interactions among duplex-DNA segments, such as DNA supercoiling, DNA looping, or segment-segment interactions promoted by polycations and other DNA-condensing agents (30–32). There is little information regarding endogenous knotting of DNA *in vivo*, although recent studies in yeast suggest that there can be low steady-state levels of knots in intracellular chromatin (33). If such knots exist *in vivo*, there must be mechanisms to efficiently resolve such topological entanglements, which are a potential death sentence for the cell (2, 34–36).

## COMPUTATIONAL METHODS

### DNA Model and Simulation Procedure

Following previous studies (18, 19, 21, 37) circular duplex DNA is modeled as a discrete semi-flexible chain with *N* extensible segments of mean length *b*_0_ = 10 nm, corresponding to a total chain length of *L* = *Nb*_0_; in this work we use *N* = 200 corresponding to 6-kbp DNA (each segment has approximately 30 bps). The potential energy of a chain conformation is given by

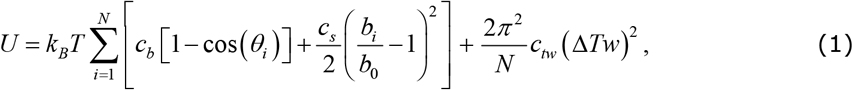

where *T* = 300 K is the temperature and *k_B_* is Boltzmann’s constant. *θ_i_* is the bending angle between successive segments *i* and *i* +1, *b_i_* is the length of segment *i*, and Δ*Tw* is the double-helical twist relative to relaxed DNA. During a Monte Carlo simulation, the value of Δ*Tw* was calculated for each chain conformation using White’s equation Δ*Tw* = Δ*Lk* − *Wr* where *Wr* is the writhe of the chain conformation and Δ*Lk* was fixed during the simulation. The bending energy constant *c_b_* is chosen such that the persistence length *P* of the chain is equal to 5 segments, i.e., *P* = 5*b*_0_ = 50 nm, resulting in *c_b_* = 5.5157 (21). The stretching energy constant is given by *c_s_* = *K_s_b*_0_/(*k_B_T*) where *K_s_* is the stretch modulus of DNA; using the approximate value *K_s_* = 1000 pN for B-form DNA under physiological conditions (38) results in *c_s_* = 2500. The twisting energy constant is given by *c_tw_* = *C*/(*b*_0_*k_B_T*) where *C* is the torsional rigidity constant of DNA; using *C* = 3×10^−19^erg·cm for B-form DNA (37) results in *c_tw_* = 7.243. Excluded-volume and electrostatic interactions between DNA segments are modeled by an effective hard-cylinder diameter *d* = 5 nm, corresponding to an ionic strength of 150 mM (39).

Equilibrium ensembles of chains with fixed knot type *K* and linking number difference Δ*Lk* were generated by Monte Carlo (MC) simulation. In our procedure, chain conformations evolved by crankshaft rotations and stretching moves of sub-chains (21), and sub-chain translations, or reptations, along the local chain axis; the purpose of reptation moves was to increase the probability of extrusion and resorption of superhelix branches (37). Trial conformations were accepted with probability *P_accept_* = min[exp[−((*U_trial_* − *U_current_*)/(*k_B_T*)),1] according to the Metropolis criterion, where *U_trial_* and *U_current_* are the potential energies of trial and current conformations, respectively, according to Equation (1). Excluded-volume interactions and preservation of knot type *K* were implemented by rejecting any trial conformation in which chain segments overlapped or which resulted in a change of *K*. Knot types *K* of current and trial conformations were determined by calculating the Alexander polynomial Δ(*t*) for *t* = −1.1 and the HOMFLY polynomial *P*(*a,z*) (40). Averages for given knot type *K* and linking number Δ*Lk* were calculated from ensembles containing 10^6^ saved conformations for the unknot 0.1 and the trefoil knot 3.1, and 5×10^5^ saved conformations for all other knot types *K*. The simulation period between saved conformations entering these ensembles was 1000 MC moves.

### Model of Type-II Enzymes

DNA-bound type-II enzymes with hairpin and straight G segments were modeled by selecting four or two contiguous chain segments, respectively, whose local geometry during a trial move conformed to specific criteria (Figure 2). A hairpin G segment formed two sides of an equilateral triangle with side lengths 2*b*_0_ = 20 nm, corresponding to a 120° bend. A putative T segment was considered to be juxtaposed with the G segment if it passed through the triangle in such a way that none of the chain segments overlapped, i.e., excluded-volume interactions were preserved for the enzyme (Figure 2A). For a straight G segment, a potential T segment was considered to be juxtaposed if it passed through an equilateral triangle with one side formed by the straight G segment of length 2*b*_0_ = 20 nm (Figure 2B). The orientation of this triangle about the center axis of the chain was chosen randomly for each trial conformation. Again, excluded-volume interactions of chain segments were preserved.

**Figure 2.**
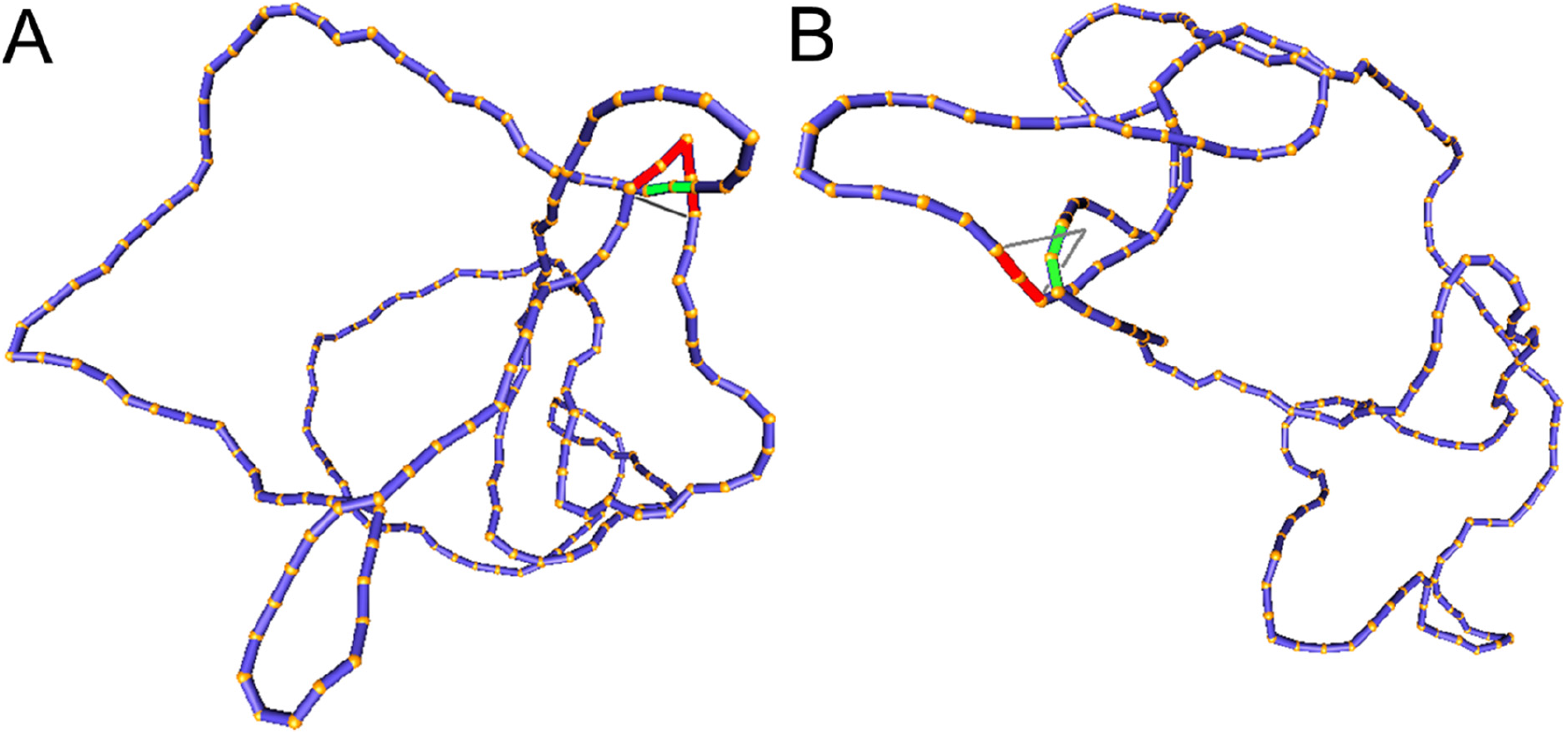
Simulation snapshots of the left-handed trefoil knot 3.1^−^ with (**A**) hairpin-like G segment and (**B**) straight G segment (59). The conformations shown correspond to states in which a T segment (green) is properly juxtaposed with the G segment (red) to initiate strand passage. Deformed chains used to determine knot type *K*’ and linking number Δ*Lk*’ of the chain conformation after strand passage are indicated by grey lines.

### Juxtaposition Probabilities and Transition Rates

Strand passages by type-II enzymes generate transitions from topological states *a* = (*K*, Δ*Lk*) to states *b* = (*K*’, Δ*Lk*’) with Δ*Lk*’ = Δ*Lk* ± 2. The associated transition rates *k_ab_* are assumed to be of the form

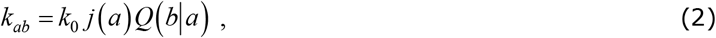

where *k*_0_ is a constant which depends on enzyme activity and concentration, but is independent of the topological states *a*, *b* of the DNA (9, 15). *j*(*a*) is the juxtaposition frequency of the enzyme in state *a*, corresponding to the fraction of DNA conformations in state *a* in which a potential ⊤ segment is properly juxtaposed with the G segment as described above. *Q*(*b*|*a*) is the conditional probability that strand passage from a juxtaposed conformation in state *a* results in state *b*. The state *b* = (*K*’,Δ*Lk*’) of the chain that would result from state *a* = (*K*, Δ*Lk*) by passage of the ⊤ segment through the G segment was determined by considering local deformations of the chain as follows (Figure 2). The knot type *K*’ was determined by calculating the Alexander and HOMFLY polynomials of the deformed chain (40). Δ*Lk*’ was determined by calculating the writhe *Wr*’ of the deformed chain and assuming that strand passage leaves the twist Δ*Tw* nearly unchanged; applying White’s equation Δ*Lk* = Δ*Tw* + *Wr* to original and deformed chain, and using Δ*Tw*’ = Δ*Tw*, then gives Δ*Lk*’–Δ*Lk* = *Wr*’–*Wr*. Using the fact that the change of Δ*Lk* occurs strictly in steps of ±2 allowed us to determine the sign of the change of Δ*Lk* by the corresponding change of the writhe *Wr*, which was always close to ±2 in our simulations. Thus, both *j*(*a*) and *Q*(*b*|*a*) can be determined by MC simulations of equilibrium ensembles of chains in a fixed topological state *a*. The validity of this approach is based on the assumption that the reaction is not diffusion limited, which implies that the probability *j*(*a*) of finding a potential T segment properly juxtaposed with the G-segment is equal to the equilibrium probability of this juxtaposed conformation in the absence of strand passage (9).

### Master Equation and Non-Equilibrium Steady States

Consider an ensemble of circular duplex DNA molecules acted on by type-II enzymes in the presence of ATP. The rates *k_ab_* for transitions from topological states *a* = (*K*,Δ*Lk*) to states *b* = (*K*′,Δ*Lk*′) induced by the enzyme are given by Equation (2). The probability *P*(*a,t*) to find a given DNA molecule in topological state *a* at time *t* obeys the master equation

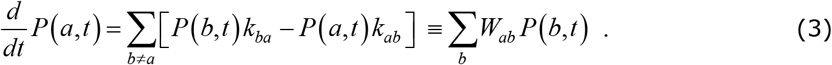

We consider here the situation where the probabilities *P*(*a, t*) are stationary, i.e., time-independent, for all topological states *a*, corresponding to non-equilibrium steady states (NESS). The stationary NESS probabilities *P*\*(*a*) were deduced from the eigenvector with eigenvalue 0 of the transition matrix *W_ab_* in Equation (3) and using the normalization condition *Σ_a_P*\*(*a*) = 1 (the star symbol for *P*\*(*a*) is used to distinguish NESS probabilities from the equilibrium probabilities *P*(*a*) obtained in ESP ensembles). Stationary NESS probability currents from topological states *a* to states *b* are found from *i_ab_* = *P*\*(*a*)*k_ab_* − *P*\*(*b*)*k_ba_*. Note that at thermal equilibrium the detailed balance condition implies *i_ab_* = 0; conversely, in our study, NESS with appreciable probability currents *i_ab_* were generated by continuously delivering a complex topology, e.g., knot type *K* = 10.139^−^ with Δ*Lk* = −12, to the ensemble by introducing a source rate *k_S_*(*a,b*) with origin *a* = (0.1,0) (the unknot with Δ*Lk* = 0) and source state *b* = (10.139^−^, −12).

### Universal NESS Probabilities and Probability Currents

For nonzero source rates *k_S_* the NESS probabilities *P*\*(*a*) and probability currents *i_ab_* depend on enzyme properties such as intrinsic rate and concentration in terms of the constant *k*_0_ in Equation (2). In order to obtain results independent of such largely unknown details (in this sense “universal”) we define normalized transition rates as

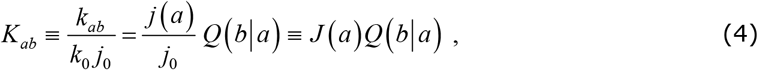

where *j*_0_ is the juxtaposition frequency in a reference state, which we choose as (0.1,0) (the unknot with Δ*Lk* = 0). The normalization factor *k*_0_*j*_0_ =Σ_*b*_*k*(0.1,0;*b*) in Equation (4), with *k*(*a;b*) = *k_ab_* from Equation (2), corresponds to the total rate of enzyme reaction in the reference state *a* = (0.1,0). The normalized juxtaposition frequency *J*(*a*) = *j*(*a*)/*j*_0_ in Equation (4) is the ratio of the actual juxtaposition frequency *j*(*a*) in state *a* and the juxtaposition frequency *j*_0_ in the reference state, where the unknown constant *k*_0_ drops out. Universal NESS probabilities *P*\*(*a*) as a function of the parameter *κ* = *k_s_*/(*k*_0_*j*_0_) were calculated using the normalized rates *K_ab_* in Equation (4) as described above, and universal NESS probability currents are obtained as *I_ab_* = *P*\*(*a*)*K_ab_* – *P*\* (*b*)*K_ba_*. The universal NESS probabilities *P*\*(*a*) and probability currents *I_ab_* as functions of the parameter *κ* are expected to depend only on geometric properties of the enzyme, such as the bend angle of the G segment. Thus these quantities are independent of properties that do not involve the particular topological state of the DNA, for example the overall size of the enzyme (as long as it is much smaller than the DNA) and the precise form of the interaction potential between the G and ⊤ segments. We verified by our simulations that *P*\*(*a*) and *I_ab_* are indeed universal functions of the parameter *κ* by showing that *P*\* (*a*) and *I_ab_* remained unchanged when altering the interaction between G and ⊤ segments (Supplementary Figure S6). This test also provided an internal control for the validity of our computational approach.

## RESULTS

### Equilibrium Distribution and Free-energy Landscape

As outlined in the Introduction, the topological distribution at thermal equilibrium provides a reference state necessary to understand ATP-driven type-II enzyme action that results in topology simplification beyond equilibrium. The equilibrium ensemble is characterized by the joint probability distribution *P*(*K*, Δ*Lk*), corresponding to an equilibrium segment-passage (ESP) ensemble of phantom chains, and distributions derived therefrom (18, 19). In particular, Podtelezhnikov *et al*. found that *P*(*K*|Δ*Lk*), the conditional distribution of *K* for given Δ*Lk*, is dominated by only a few knots *K* for any fixed value of Δ*Lk*; moreover, the dominating knots except for the unknot were all chiral (18). Later, Burnier *et al*. pointed out that for chiral knots *K* the level of supercoiling is characterized by the quantity Δ*Lke* = Δ*Lk* – 〈*Wr*〉(*K*,nicked) rather than Δ*Lk*, where 〈*Wr*〉(*K*,nicked) is the signed, nonzero mean value of the 3D writhe for a torsionally unconstrained (nicked) DNA molecule with chiral knot type *K* (19). This result can be easily understood by taking the average of White’s equation for fixed Δ*Lk*, i.e., Δ*Lk* = (Δ*Tw*) + 〈*Wr*〉 : for a torsionally relaxed, i.e., not supercoiled, chain one has 〈Δ*Tw*〉 = 0 and 〈*Wr*〉 = 〈*Wr*〉(*K*,nicked), thus Δ*Lk* = 〈*Wr*〉(*K*,nicked) and Δ*Lke* = 0 (Figure 3). Burnier *et al*. found that the conditional distribution *P*(Δ*Lke*) is dominated by the unknot for *any* fixed value of Δ*Lke*; moreover, *P*(*K*|Δ*Lke*) decreases with increasing −Δ*Lke* for any knot *K*, implying that increasing levels of supercoiling favor unknotting (19).

**Figure 3.**
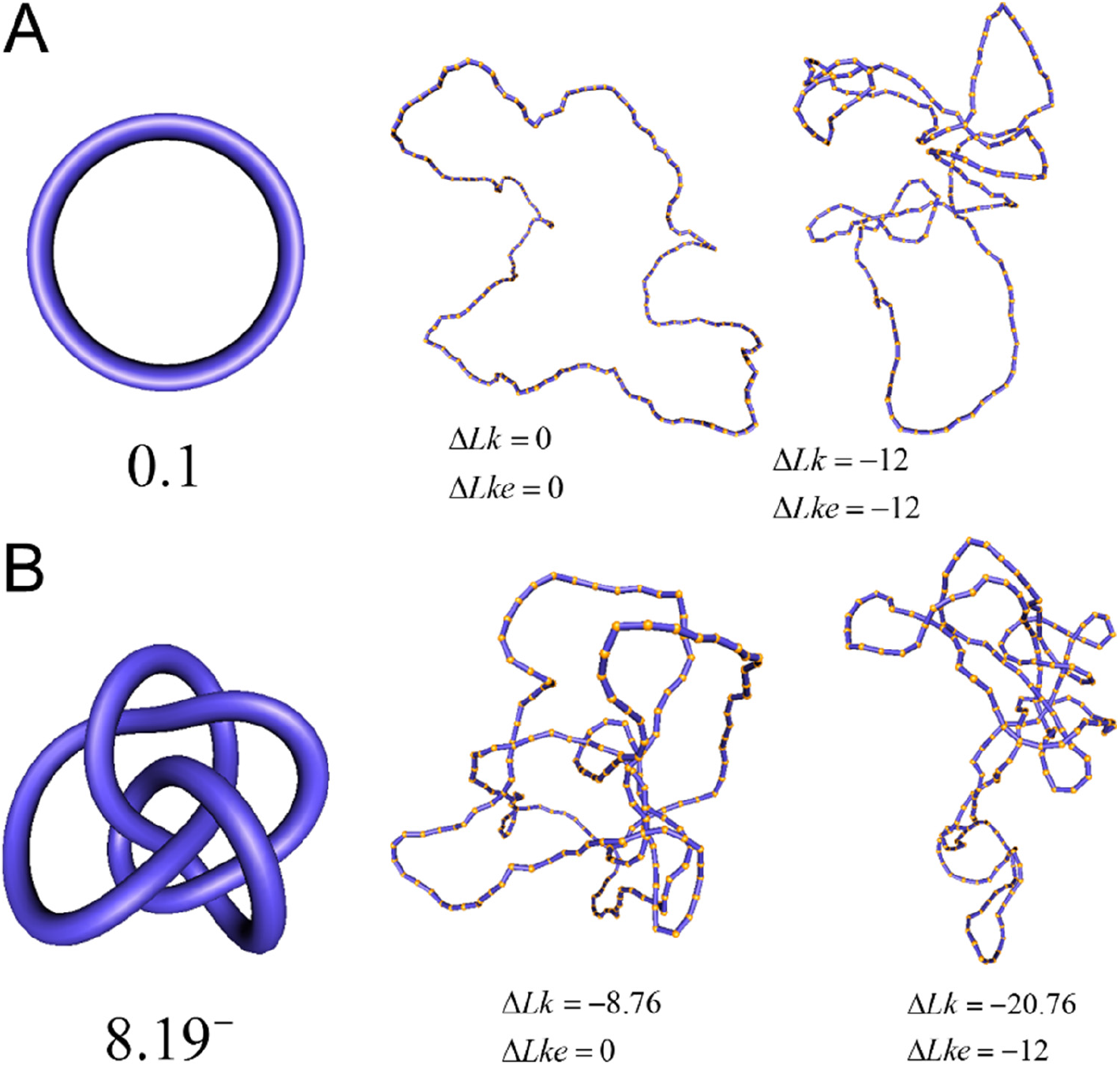
Standard forms of (**A**) the unknot 0.1 and (**B**) the left-handed knot 8.19^−^, and simulation snapshots of 6-kbp DNAs of these knots with values of Δ*Lk* and Δ*Lke* = Δ*Lk* − 〈*Wr*〉 (*K*,nicked) as shown (59). The mean writhe 〈*Wr*〉(*K*,nicked) of torsionally relaxed (nicked) DNA is 0 for *K* = 0.1 and −8.76 for *K* = 8.19^−^. The states with Δ*Lke* = 0 appear relaxed, whereas for Δ*Lke* = −12 supercoiling is present.

We first verified that our calculation reproduces the behavior of the equilibrium distribution *P*(*K*|Δ*Lk*) found earlier (see Figure 4 in reference (18) and Figure 2A in reference (19)). For our 6-kbp DNAs we indeed find that for any fixed, small value of Δ*Lk* only a few knot types *K* dominate the distribution. However, for -Δ*Lk* > 18, corresponding to superhelix density −*σ* = Δ*Lk*/*Lk*_0_ >0.0315 for 6-kbp DNAs, the distribution rapidly becomes degenerate and many different knot types *K* contribute to *P*(*K*|Δ*Lk*) (Supplementary Figure S2). This value of *σ* is closely similar to the *in-vivo* level of unconstrained supercoiling in prokaryotes (41, 42). If conditions inside the cell increase the level of unconstrained supercoiling beyond this |*σ*| value, the resulting distribution of knot types would be expected to become highly degenerate.

**Figure 4.**
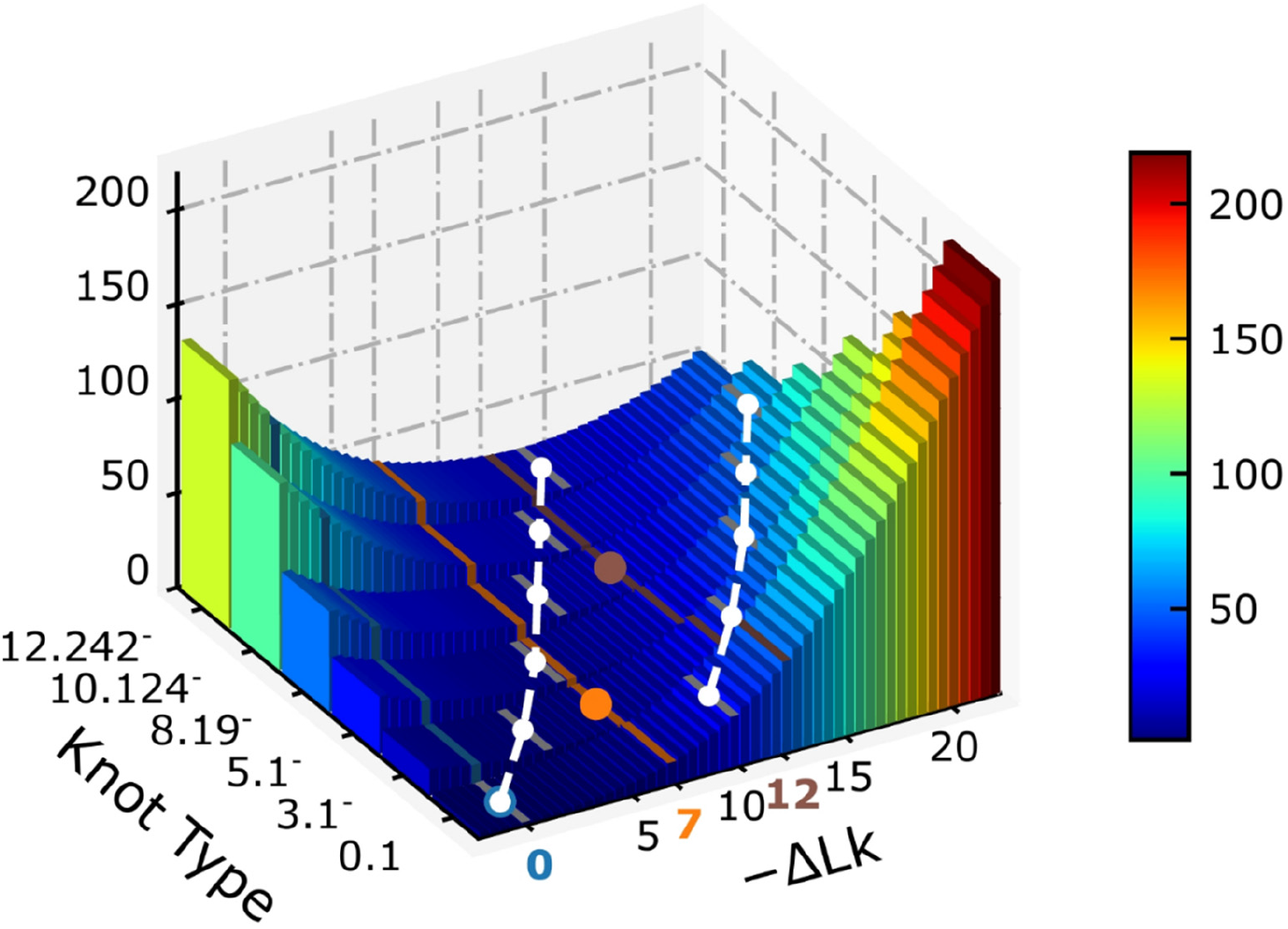
Free energy landscape *F*(*K*,Δ*Lk*) = −*k_B_T*ln[*P*(*K*,Δ*Lk*) in units of *k_B_T* for 6-kbp DNAs, where *P*(*K*,Δ*Lk*) is the joint probability distribution of *K* and Δ*Lk*. Along sections with fixed −Δ*Lk* > 5.5, *F*(*K*, Δ*Lk*) is minimum for nontrivial knots, i.e., knots different from the unknot 0.1. This is indicated for −Δ*Lk* = 7 (*F* minimum for *K* = 3.1^−^, orange line/dot) and −Δ*Lk* = 12 (*F* minimum for *K* = 8.19^−^, brown line/dot). The white curves are sections for fixed values 0, −10 of the degree of supercoiling Δ*Lke* = Δ*Lk* − 〈*Wr*〉 (*K*,nicked) where 〈*Wr*〉 (*K*,nicked) is the mean writhe for torsionally relaxed (nicked) DNA with knot type *K*. Along these sections, *F* is always minimum for the unknot 0.1 and the corresponding free energy gradient towards 0.1 becomes steeper with increasing −Δ*Lke*.

Next, in order to understand the most-probable relaxation path of a given DNA knot *K* with linking number Δ*Lk* by a topoisomerase that is driven only by the topological free-energy gradient, we calculated the free energy landscape *F*(*K*, Δ*Lk*) = −*k_B_T* ln *P*(*K*, Δ*Lk*) including all knot types *K* which dominate the distribution *P*(*K*| Δ*Lk*) and have 12 or fewer crossings (Figure 4). (See Supplementary Data, Section S1, Figure S1 and Table S1 for details regarding the calculation of *P*(*K*,Δ*Lk*)). The free-energy landscape also explains the apparent contradiction between results for *P*(*K*|Δ*Lk*) and *P*(Δ*Lke*) obtained in references (18) and (19), respectively, by noting that distributions for fixed Δ*Lk* or Δ*Lke* merely correspond to different sections of the same free energy landscape *F*(*K*,Δ*Lk*) (Figure 4): along sections with fixed −Δ*Lk* >5.5, the minimum value of *F*(*K*,Δ*Lk*) corresponds to the chiral knot 3.1^−^, whereas along sections with fixed Δ*Lke* = Δ*Lk* − 〈*Wr*〉 (*K*,nicked),the minimum in *F* always coincides with the unknot 0.1. The corresponding free-energy gradient towards 0.1 is steeper for increasing −Δ*Lke*, in agreement with earlier results (18, 19).

### Steady-State Knot Distributions in Supercoiled DNA and Topology Simplification

In addressing the influence of DNA supercoiling on the unknotting efficiency of type-II enzymes, we first consider an ensemble of supercoiled 6-kbp circular duplex DNAs in the presence of type-II topoisomerase and ATP without additional components. Each round of type-II enzyme action converts a DNA substrate in the state (*K*,Δ*Lk*) to a product state (*K*’,Δ*Lk*’) where Δ*Lk*’ = Δ*Lk*± 2 and *K*’ is a knot that can be obtained from *K* by one intersegmental passage (43). Figure 5 shows steady-state fractions *P*\*(*K*, Δ*Lk*) of the unknot 0.1 and stereoisomers 3.1+ and 3.1^−^ of the trefoil knot for type-II enzymes modeled in terms of hairpin-like and straight G segments, respectively. Knots with more than 3 crossings occurred with low frequency and were omitted from Figure 5 for simplicity. As a comparison we also show the equilibrium probabilities *P*(*K*,Δ*Lk*) corresponding to ESP ensembles.

**Figure 5.**
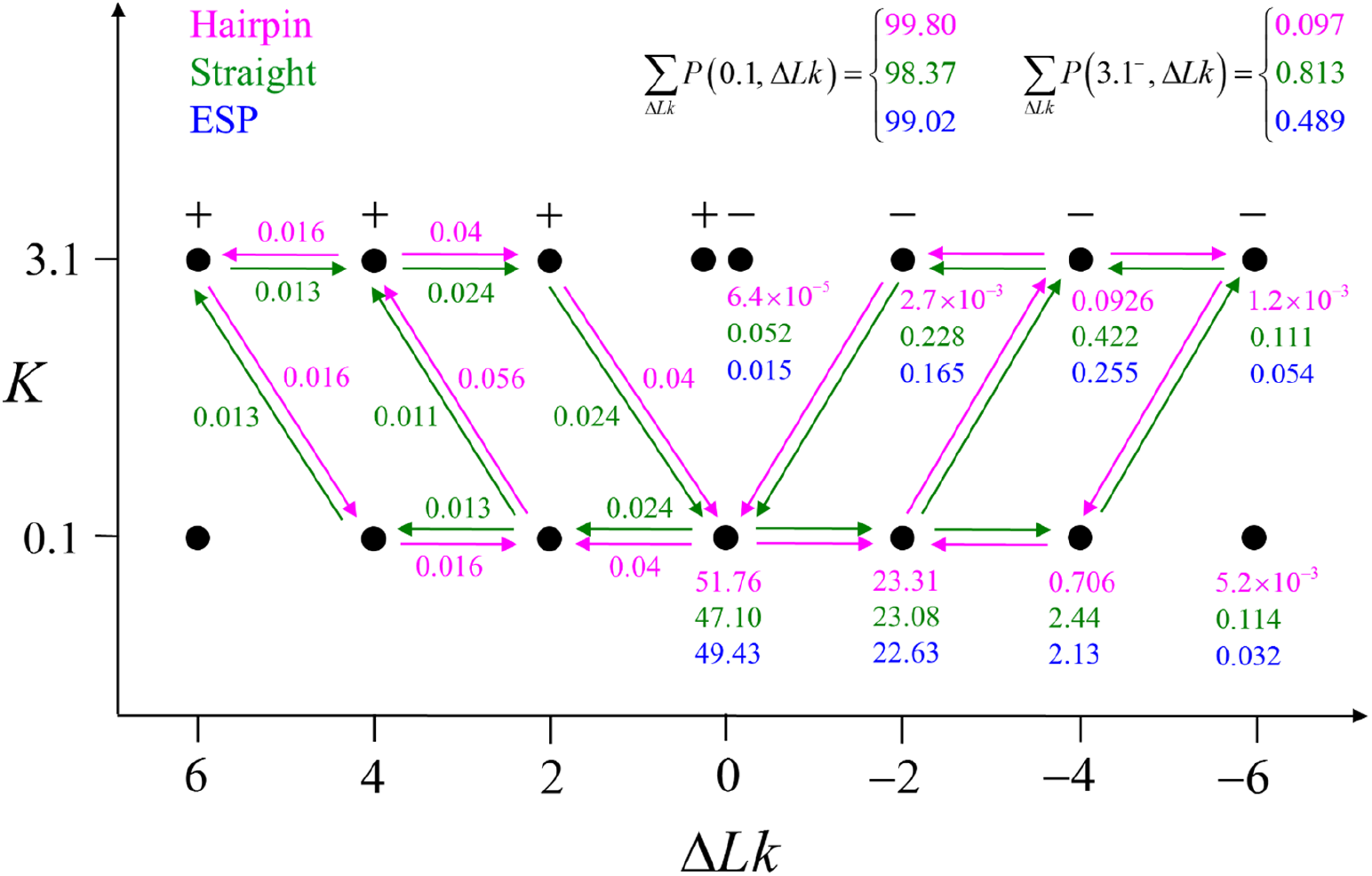
Steady-state fractions *P*\*(*K*,Δ*Lk*) for the unknot 0.1 and isoforms 3.1+ and 3.1^−^ of the trefoil knot in the presence of type-II enzymes modeled with hairpin-like and straight G segments, respectively. Also shown are the corresponding equilibrium fractions *P*(*K*, Δ*Lk*) for comparison. The group of numbers shown for each knot *K* are steady-state fractions *P*\*(*K*,Δ*Lk*), in percent, for DNAs with hairpin-like G segments (upper entries, magenta), straight G segments (middle entries, green), and equilibrium fractions *P*(*K*, Δ*Lk*) corresponding to ESP ensembles (lower entries, blue). The fractions for right-handed isoforms of a chiral knot are the same as for left-handed isoforms by symmetry. The upper panel displays sums of *P*\*(*K*,Δ*Lk*) over the Δ*Lk* - values shown in the figure. The arrows and associated numbers indicate residual probability currents *I*(*K*, *K*’) for enzymes with hairpin (magenta) and straight G segments (green).

Figure 5 shows that the steady-state fraction of 3.1 knots is reduced for enzymes with hairpin G segments compared to enzymes with straight G segments, consistent with results obtained earlier for torsionally unconstrained (nicked) chains (9). For the sums *P*\*(0.1) = Σ_Δ*LK*_*P*\*(0.1,Δ*Lk*) and *P*\*(3.1^−^) = Σ_Δ*Lk*_*P*\*(3.1^−^, Δ*Lk*), corresponding to steady-state probabilities of knots 0.1 and 3.1^−^ for nicked chains, we find *P*\* (3.1^−^)/*P*\*(0.1) = 9.7 ×10^−4^ (hairpin G segment) and *P*\*(3.1^−^)/*P*\*(0.1) = 0.0079 (straight G segment). This corresponds to a reduction by a factor of about 8 (compare column *C_k_*/*C_u_* in Table 1 in reference (9), where both isoforms 3.1^−^ and 3.1+ were included in the statistics of the trefoil knot 3.1 for nicked 7-kbp DNAs). The difference in reduction factors of 14 in reference (9) and 8 in our study may be explained by the fact that the hairpin-like G segment considered in (9) had an overall 180°-bend compared to a smaller 120° - bend in our model (see Computational Methods).

However, for fixed values of Δ*Lk* the reduction factor depends strongly on the value of Δ*Lk*; for example, for Δ*Lk* = −4 the reduction factor is only 1.5 whereas for Δ*Lk* = 0 it is 61 (Figure 5). The dependence of the reduction factor on Δ*Lk* is related to the fact that the free-energy gradient depends on the relevant section of the free-energy landscape *F*(*K*,Δ*Lk*) : the gradient toward 0.1 is much steeper for fixed Δ*Lk* = 0 than for Δ*Lk* = −4 (Figure 4). We also found that the steady-state fractions *P*\*(*K*,Δ*Lk*) for knots *K* different from the unknot are slightly larger for type-II enzymes with straight segment than the corresponding equilibrium probabilities *P*(*K*, Δ*Lk*) (Figure 5), again in agreement with results obtained previously for nicked chains (compare reference (9), Table 1). Interestingly, for supercoiled DNA, residual cycle-probability currents appear. Such cyclic probability currents occur because type-II enzymes drive the reaction away from thermal equilibrium so that detailed balance between directed fluxes (*K*, Δ*Lk*)→(*K*’, Δ*Lk*’) and (*K*’,Δ*Lk*’)→(*K*,Δ*Lk*) is violated in general. However, it is not clear whether these residual probability currents have any significance regarding the unknotting efficiency of type-II enzymes.

Apart from DNA unknotting, another aspect of DNA-topology simplification by type-II enzymes is a reduction of the degree of supercoiling, which translates into a narrower Δ*Lk* -distribution about its mean value 〈*Wr*〉(*K*,nicked) for a given knot type *K*. A metric used to quantify this type of topology simplification is the topology simplification factor *TSF* = s.d.(Δ*Lk*,topo II)/s.d.(Δ*Lk*,topo I) where s.d.(Δ*Lk*,topo II) is the standard deviation of Δ*Lk* in the presence of type-II enzyme and ATP, and s.d.(Δ*Lk*,topo I) is the standard deviation of Δ*Lk* in the presence of type-I enzyme. The latter does not consume energy from ATP hydrolysis and thus generates the Δ*Lk* distribution corresponding to an ESP ensemble at thermal equilibrium. In reference (8) the variance of the Δ*Lk* -distribution was measured for the nicked unknot form of 7-kbp pAB4 DNA in the presence of *E. coli* topoisomerase IV and ATP and gave the result 〈Δ*Lk*〉= 1.7 compared with the equilibrium value 3.1, which yields *TSF* = 0.74. We studied the narrowing of the Δ*Lk* distribution for the unknot 0.1 in the presence of type-II enzymes modeled with the hairpin-like G segment and compared the standard deviations of the steady-state distribution *P*\*(Δ*LK*|0.1) and the equilibrium distribution *P*(Δ*Lk*|0.1) (Supplementary Data, Section S3 and Figure S3). Note that the distributions *P*\*(Δ*Lk*|0.1) for even and odd values of Δ*Lk* are disjunct because type-II enzymes change Δ*Lk* in steps of 2; conversely, type-I enzymes change Δ*Lk* in steps of 1. We thus find *TSF* = 0.91 for Δ*Lk* even and *TSF* = 0.86 for Δ*Lk* odd, in reasonable agreement with the experimental result (8) (Supplementary Data, Section S3).

### Pathways of Topology Simplification in Knotted, Supercoiled DNA

For DNAs in the size range considered here and in the absence of a process that actively delivers a complex knot type to the ensemble of DNAs, the equilibrium probabilities *P*(*K*,Δ*Lk*) are very small for any knot *K* different from the unknot. In the presence of a type-II enzymes these probabilities are reduced even further. Thus, for DNA molecules a few kbp in length practically no knotted DNAs appear even in the absence of type-II enzymes. However, a typical situation *in vivo* is that some biological process is present that actively generates knotted DNAs, and type-II enzymes are essentially needed to remove these knots. To address this biologically relevant situation, we now assume the presence of an extraneous process that continuously delivers DNA molecules in a complex source state *a_S_* = (*K_S_*, Δ*Lk_S_*). Specifically, we assume that a process is present in the ensemble of 6-kbp duplex DNAs that continuously converts unknotted DNAs with Δ*Lk* = 0 to DNAs forming the knot 10.139^−^ with linking number Δ*Lk* = −12 at constant rate *k_S_*. The knot 10.139^−^ contributes notably to the distribution *P*(*K*|Δ*Lk*) at Δ*Lk* = −12 (Supplementary Figure S2) and is chosen here to illustrate the pathway of topology simplification by type-II topoisomerase given an initial complex topological state.

The DNA molecules delivered in the source state *a_S_* = (10.139^−^,−12) by the extraneous process are converted by type-II enzyme strand passages to simpler topological forms in a stepwise manner, resulting in a pathway of intermediate topological states. As discussed in the previous section, each round of type-II enzyme action converts a DNA substrate in the state (*K*,Δ*Lk*) to a product state (*K*’,Δ*Lk*’) where Δ*Lk*’ = Δ*Lk*±2 and *K*’ is a knot that can be obtained from the knot *K* by one intersegmental passage (43). Eventually the DNAs are converted back to the originating state (0.1,0), i.e., the unknot with Δ*Lk* = 0. The latter is then converted again to molecules in the source state *a_S_* = (10.139^−^,−12) by the extraneous process, resulting in a continuous cycle. The cyclic process is characterized by non-equilibrium steady state (NESS) probabilities *P*\* (*a*) for DNAs in topological states *a* = (*K*,Δ*Lk*), and probability currents *I_ab_* for transitions from states *a* = (*K*,Δ*Lk*) to *b* = (*K*’,Δ*Lk*’). The NESS probabilities *P*\*(*a*) are appreciable for the source state *a_S_* and all intermediate states *a* along the pathway of topology simplification by topoisomerase-II action.

Figure 6 shows resulting pathways of topology simplification for type-II enzymes modeled by a hairpin-like (Figure 6A) and straight G segments (Figure 6B), respectively. Steady-state probabilities *P*\*(*a*), in percent, are shown next to each state *a* = (*K*, Δ*Lk*), and probability currents *I_ab_* are given by numbers next to the arrows. The source probability current associated with the external process that converts DNAs in the originating state (0.1,0) to the source state *a_S_* = (10.139^−^,−12) is given by *I_S_* = *P*\* (0.1,0)*κ* with *κ* = *k_S_*/(*k*_0_*j*_0_) (see Computational Methods). Only probability currents *I_ab_* with *I_ab_*/*I_S_* > 0.05 are shown, where dominant currents with 0.05 < *I_ab_*/*I_S_* > 0.1 are shown as dark blue arrows and subdominant currents with 0.05 < *I_ab_*/*I_S_* < 0.1 are shown as light blue arrows. Empty circles indicate states (*K*, Δ*Lk*) for which Δ*Lke* = Δ*Lk* − 〈*Wr*〉(*K*,nicked) = 0, corresponding to torsionally relaxed chains (cf. white curve on the left in Figure 4). It is apparent that the pathways shown on the left sides in Figures 6A, 6B closely follow the path Δ*Lke* = 0. Interestingly, only a small number of intermediates contribute to the pathways although there exist about 250 different knot types with 10 or fewer crossings.

**Figure 6.**
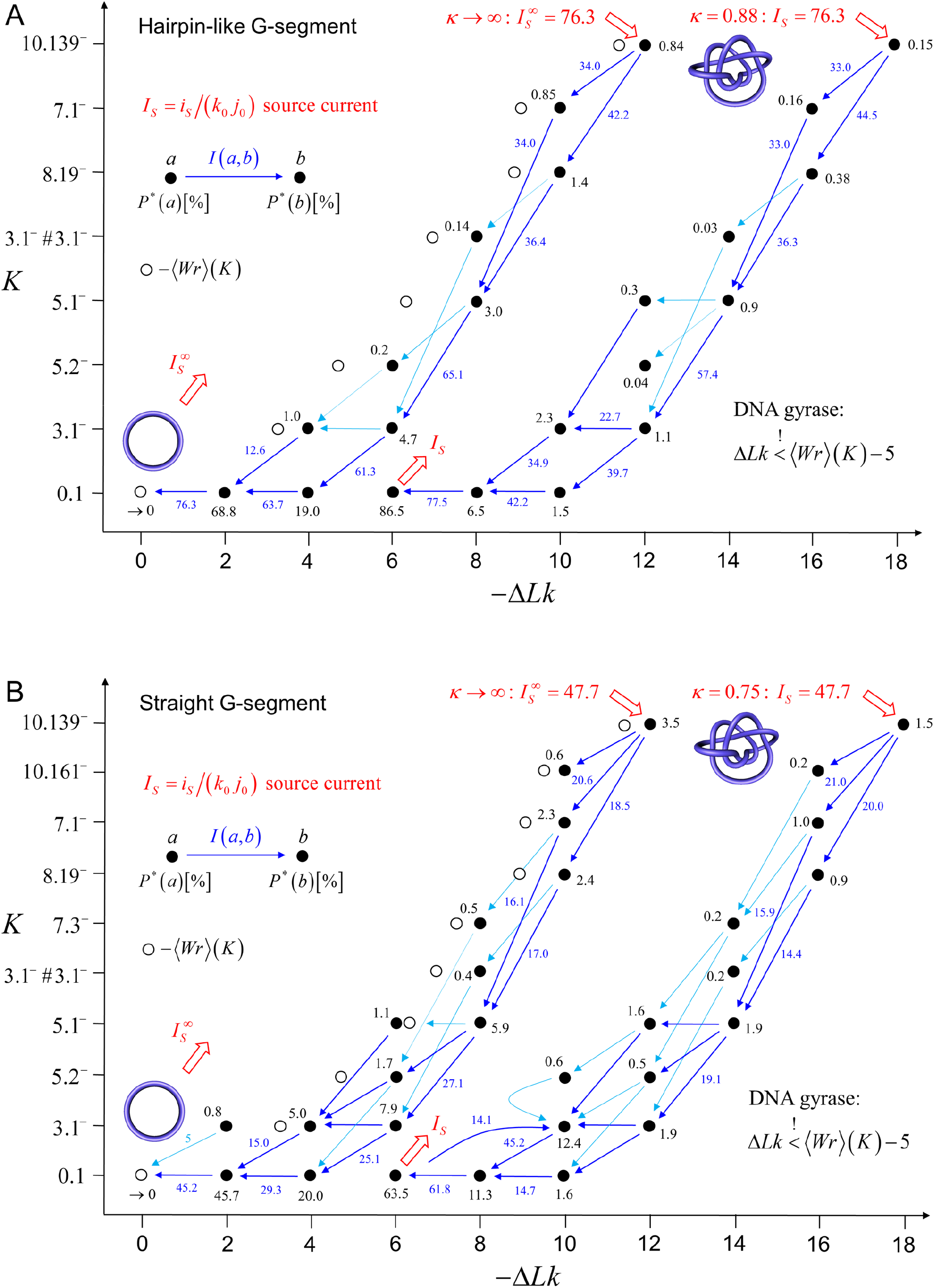
Comparison of NESS probabilities and probability currents generated by type-II topoisomerase activity with (**A**) hairpin and (**B**) straight G segment (59). In both cases, we imposed an external process that converts unknotted DNA with Δ*Lk* = 0 to DNA forming a (source) knot *K_S_* = 10.139^−^ with Δ*Lk*_S_ = −12 in the limit of large source rate *k_S_* (pathways shown on the left in (**A**) and (**B**)). Dominant probability currents with 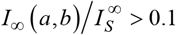 are shown as dark blue arrows and subdominant probability currents with 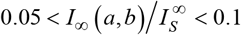 are shown as light blue arrows. Steady-state probabilities 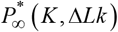, in percent, are shown next to each knot *K*. Open circles indicate positions of Δ*Lk* = 〈*Wr*〉 (*K*,nicked), i.e., Δ*Lke* = Δ*Lk* − 〈*Wr*〉 (*K*,nicked) = 0 (cf. white curve on the left in Figure 4). The pathways shown on the right in (**A**) and (**B**) show cases in which a supercoiled state is maintained by introducing the constraint Δ*Lke* < −5. In these cases, we assumed the presence of an external process that converts unknotted DNA with Δ*Lk* = −6 to DNA forming a source knot *K_S_* = 10.139^−^ with Δ*Lk_S_* =−18, and the source rate *k_S_* was adjusted to obtain the same source probability current *I_S_* = 76.3 as for the pathway shown on the left to facilitate comparison.

For the pathways shown on the left sides in Figures 6A, 6B we consider the limit of a large source rate *k_S_* for the external process. In this limit, the originating state (0.1,0) is depleted by the external process, which implies that the steady-state probability of the originating state vanishes as *P*\* (0.1,0) ~ 1/*k_S_*. For all other states *a* the steady-state probabilities approach finite values 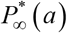 in the limit of a large source rate *k_S_*. Likewise, all probability currents *I_ab_* approach finite values 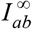 in the limit of large source rate *k_S_*, including the source probability current *I_S_*. Therefore, the values of the steady-state probabilities *P*\*(*a*) and probability currents *I_ab_* in Figures 6A, 6B are universal in the sense that they are independent of the precise value of the source rate *k_S_* as long as *k_S_* is large enough. The full dependence of *P*\*(*a*) and *I_ab_* on the parameter *κ* = *k_S_*/(*k*_0_*j*_0_) is shown in Supplementary Figures S4, S5.

In many biological systems a finite amount of supercoiling is maintained. For example, for bacterial cells the torsional tension is maintained by a homeostatic mechanism involving topoisomerase I and DNA gyrase (44, 45). To study this situation, on the right sides in Figures 6A, 6B we show pathways of topology simplification for the case that a state of finite DNA supercoiling is maintained by introducing the constraint Δ*Lk* < −5. For these pathways we assume that an external process is present that continuously converts DNAs in the originating state (0.1,−6) to the source state *a_S_* = (10.139^−^,−18). The parameter *κ* = *k_S_*/(*k*_0_*j*_0_) is adjusted so as to produce the same source probability current *I_S_* as for the pathways shown on the left sides in Figure 6A, 6B to facilitate a comparison. Figure 6 reveals the dependence of the unknotting capability of a type-II enzyme on the degree of supercoiling. For the source state *a_S_* =(10.139^−^,−12) of the pathways shown on the left sides in Figures 6A, 6B we find 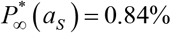 for hairpin G segment and 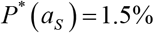 for straight G segment, corresponding to a reduction by a factor of 4.2. Conversely, for the source state *a_S_* = (10.139^−^,−18) of the pathways shown on the right sides in Figures 6A, 6B, for which the DNAs are more supercoiled, we find *P*\*(*a_S_*) = 0.15% for the hairpin G segment and *P*\*(*a_S_*) = 1.5% for the straight G segment, corresponding to a larger reduction factor of 10. Thus, supercoiling favors unknotting for the present non-equilibrium situation where a complex knot type is continuously delivered to the ensemble of DNA conformations.

Interestingly, the pathways for hairpin and straight G segments are somewhat similar. This surprising result will be explained further below in terms of juxtaposition probabilities *J*(*a*) and transition probabilities *Q*(*b*|*a*) for enzymes with hairpin and straight G segments.

### How do Type-II Enzymes with Hairpin G Segments Suppress Knotting Below Equilibrium?

As discussed in the previous section, a type-II topoisomerase with hairpin G segment reduces the steady-state fraction of complex knots below the equilibrium value relative to an enzyme with a straight G segment; moreover, the unknotting efficiency of the hairpin enzyme increases with DNA supercoiling. To better understand the origin of this effect, Figure 7 compares normalized juxtaposition probabilities *J*(*K*,Δ*Lk*) = *j*(*K*,Δ*Lk*)*j*_0_ and transition probabilities *Q*(*b*|*K*, Δ*Lk*) appearing in Equation (4) for type-II enzymes with straight (*j*_0_ = 0.0017) and hairpin G segments (*j*_0_ = 0.00013), respectively. The quantity *j*_0_ denotes the juxtaposition probability for the reference state (0.1,0) so that *J* (0.1,0) = 1 by definition (see Computational Methods). Figure 7 shows the dependence of *J* and *Q* on states (*K*, Δ*Lk*) for the knots *K* = 0.1, 3.1^−^, 8.19^−^ as a function of Δ*Lk*. 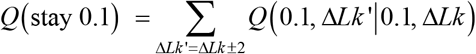 is the probability that strand passage in an unknot with linking number Δ*Lk* again results in an unknot (with Δ*Lk*’ = Δ*Lk*±2), i.e., no knotting occurs. For *K* = 3.1^−^ and 8.19^−^,

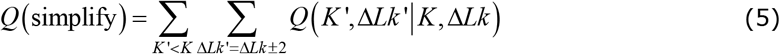

is the probability that strand passage results in unknotting, i.e., yields a knot *K*’ with a smaller number of crossings than *K* (denoted *K*’ < *K*).

**Figure 7.**
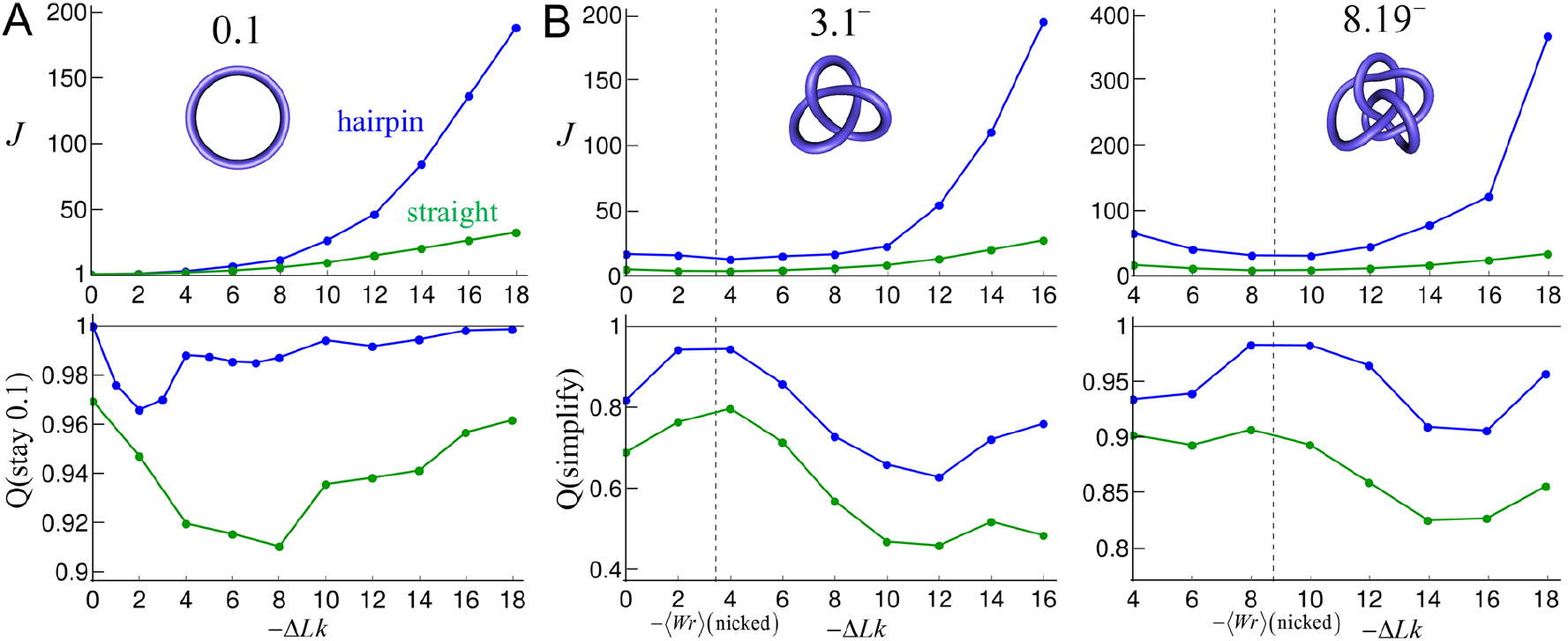
Normalized juxtaposition frequencies *J*(*K, ΔLk) = *j*(*K*, Δ*Lk*)/*j*_0_ and transition probabilities *Q*(*b*|K,Δ*Lk*) for type-II enzymes modeled by hairpin-like G segment (blue curves) and straight G segment (green curves) for knot types (**A**) 0.1 and (**B**) 3.1^−^, 8.19^−^ (59). The quantity *j*_0_ denotes the juxtaposition probability for the reference state (0.1,0), with *j*_0_ = 0.00013 for hairpin G-segment and *j*_0_ = 0.0017 for straight G-segment. Note that *J*(0.1,0) = 1 by definition. *Q*(stay 0.1) is the probability that strand passage in an unknot with linking number Δ*Lk* again results in an unknot (with Δ*Lk*’ = Δ*Lk**±2), i.e., no knotting occurs. For *K* = 3.1^−^, 8.19^−^, *Q*(simplify) is the probability that strand passage results in unknotting, i.e., in a knot *K*’ with a smaller number of crossings than *K* (see text). The vertical lines indicate values 〈*Wr*〈 (3.1^−^,nicked) = 3.433 and 〈*Wr*Ⱥ(8.19^−^,nicked) = 8.761, respectively, corresponding to Δ*Lk* - values for which the degree of supercoiling vanishes, i.e., Δ*Lke* = Δ*Lk* − 〈*Wr*〉 (*K*,nicked) = 0 (cf. Figure 3). Supercoiled chains correspond to Δ*Lk* - values to the left and right from these vertical lines.

As shown in Figure 7 (upper panels), the normalized juxtaposition probabilities *J*(*K*, Δ*Lk*) are larger for hairpin than for straight G segment, and this effect increases with the complexity of the knot *K* and with the degree of supercoiling Δ*Lke* = Δ*Lk* − 〈*Wr*〉 (*K*,nicked). The fact that *J*(*K*, Δ*Lk*) increases with knot complexity is expected because complex knots are more compact than less complex knots on average, so that more complex knots have higher probabilities of segment juxtaposition. This is consistent with the corresponding behavior of the unknot 0.1 compared with the trefoil knot 3.1^−^ for nicked DNA (9). However, for supercoiled DNA, *J*(*K*, Δ*Lk*) also increases with the degree of supercoiling Δ*Lke*, and this effect is dramatically larger for type-II enzymes with hairpin versus straight G segments. This can be qualitatively explained in terms of correlated juxtaposition of chain segments. In juxtaposed conformations of type-II enzymes with hairpin G segments, typically two crossings of the chain are made by the juxtaposed ⊤ and hairpin G segments; conversely, in juxtaposed conformations with straight G segment typically only one crossing is made by the juxtaposed ⊤ and straight G segments (Figure 8) (9). This leaves, on average, one extra crossing that has to be absorbed by the rest of the chain for the hairpin case compared with a straight G segment. The free energy *F* of unknotted supercoiled DNA increases quadratically with the superhelix density −*σ* = Δ*Lk*/*Lk*_0_, i.e., *F* ∝ (Δ*Lk*/*Lk*_0_)^2^ (see, e.g., equation (8) in (46)). Assuming that this relationship generalizes for knotted, supercoiled DNA to *F* ∝ (Δ*Lke*/*Lk*_0_)^2^ and that the extra crossing involved in the case of the hairpin G segment amounts to an increment |Δ*Lke*(hairpin)| = |Δ*Lke*(straight)| +1 in linking number that has to be absorbed by the rest of the chain, we find *F*(hairpin)−*F*(straight) ∝ (|Δ*Lke*| +1)^2^ − (Δ*Lke*)^2^ = 2 |Δ*Lke*| +1 (here Δ*Lke* = Δ*Lke*(straight)). This linear increase in free energy as a function of |Δ*Lke*| corresponds to an exponential increase in juxtaposition probability for hairpin compared to straight G segments, i.e., *J*(hairpin)/*J*(straight) ~ exp (2|Δ*Lke*|) (Figure 7, upper panel).

**Figure 8.**
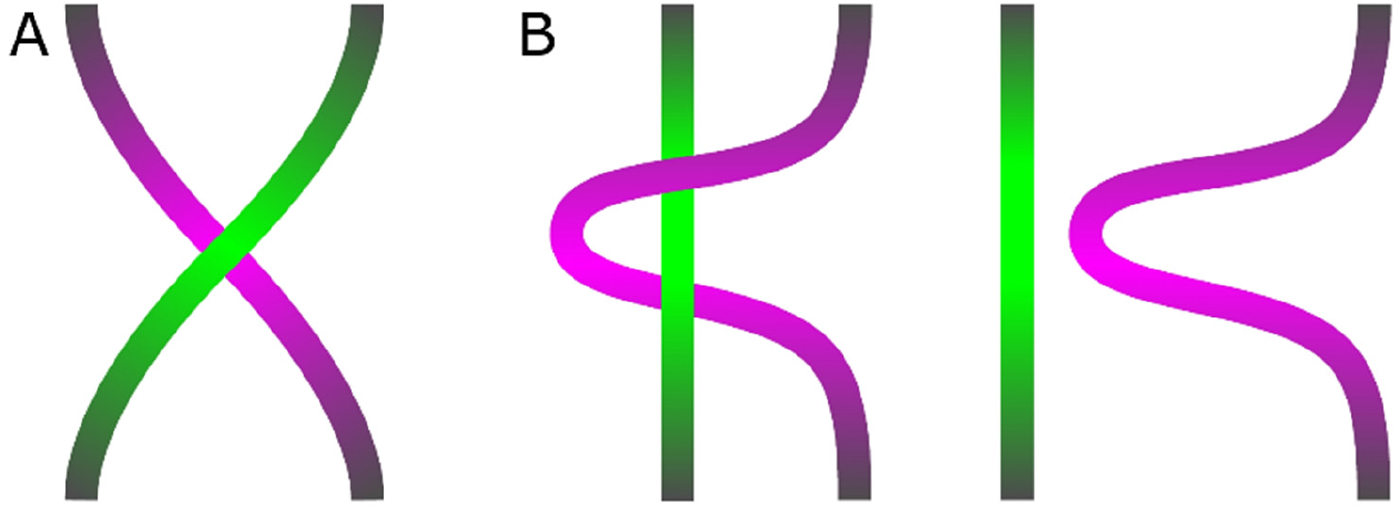
Schematic depiction of juxtaposed and passed conformations of type-II enzymes modeled by straight and hairpin G segments, respectively. The G segment is indicated by the purple portion and the ⊤ segment by the green portion of the chain. (**A**) Juxtaposed and passed conformations for straight G segment. (**B**) Juxtaposed (left) and passed (right) conformations for hairpin G segment. Note the different number of crossings made by the ⊤ and G segments.

A similar argument also explains the behavior of *Q* (stay 0.1) as a function of Δ*Lk* for hairpin compared to straight G segments (Figure 7, lower panel, left). In a conformation generated by the passage of a T segment through a hairpin G segment, corresponding to the juxtaposition of a straight segment to the outside of a hairpin, typically the passed T and hairpin G segments do not cross (Figure 8). Thus, if the passed conformation is knotted, all of the crossings of the knot have to be absorbed by the rest of the chain. Conversely, in passed conformations with straight G segments typically one crossing is made by the passed T and straight G segments, leaving one crossing less that has to be absorbed by the rest of the chain if the passed conformation is knotted (Figure 8) (9). Thus, a similar argument as above leads to *Q*(become knotted,hairpin) / *Q*(become knotted,straight) ~ exp(−2|Δ*Lk*|); see Figure 7 (lower panel, left) where *Q*(stay 0.1) = 1 − *Q*(become knotted). Interestingly, and in opposition to the behavior of *Q*(stay 0.1), the probability *Q*(simplify) that strand passage in the nontrivial knots 3.1^−^, 8.19^−^ results in unknotting is similar for hairpin and straight G segments, albeit somewhat larger for the hairpin G segment (Figure 7, lower panel middle and right). This may be explained as follows. Consider, for example, a trefoil knot which is transformed to an unknot by strand passage. Both for hairpin and straight G-segments, there is no extra crossing that needs to be absorbed by the rest of the chain in the resulting unknot after strand passage. Thus the transition probabilities *Q*(0.1, Δ*Lk* ± 2 ļ 3.1, Δ*Lk*) are expected to be similar for hairpin and straight G segments regardless of the value of Δ*Lk* (Figure 7, lower panel, middle).

Thus, the unknotting capability of type-II enzymes for complex knots is enhanced for a type-II enzyme with hairpin G segment compared to a type-II enzyme with straight G segment mainly due to a combination of two effects: 1. Enhanced juxtaposition probability *J*(*K*, Δ*Lk*) for complex knots and 2. enhanced probability *Q* (stay 0.1) for an unknot to stay unknotted after strand passage. Both these effects increase exponentially with the degree of supercoiling Δ*Lke* = Δ*Lk* − 〈*Wr*〉(*K*,nicked). Note that this effect cannot be explained alone by the free-energy landscape *F*(*K*, Δ*Lk*) (Figure 4) but is a result of the non-equilibrium dynamics associated with type-II action. Conversely, the probability *Q*(simplify) that strand passage in a complex knot results in unknotting is similar for hairpin and straight G segments, and thus does not contribute much to the unknotting capability of type-II enzymes with a hairpin versus a straight G segment. In this sense, type-II enzymes with hairpin G segments are not “smarter” than type-II enzymes with straight G segments (the latter corresponding to the equilibrium situation) but are more efficient mainly due to enhanced frequencies of juxtaposition *J*(*K*,Δ*Lk*) and probabilities *Q*(stay 0.1).

## DISCUSSION

Although much attention, experimentally and theoretically, has been devoted to understanding the action of type-II topoisomerases on unknotted, supercoiled DNA and relaxed or nicked, knotted DNA, respectively, there has been little examination of type-II enzyme activity on DNAs that are both knotted and supercoiled. Whereas negative (-) supercoiling is acknowledged to be essential for normal transactions involving DNA in living systems, unresolved knotting of a genome is generally believed to be fatal to the cell (2, 34–36). The question of how homeostatic mechanisms properly regulate supercoiling and completely eliminate knots at the same time hinges on detailed understanding of the respective rates for linking-number changes versus unknotting. Toward that end we have developed a model based on a network of topological states (*K*,Δ*Lk*) of circular DNAs with knot type *K* and linking-number difference Δ*Lk* in which the dynamics of transitions between states (*K*,Δ*Lk*) mediated by type-II enzymes is described by a chemical master equation. For the special case that the non-equilibrium fractions of states (*K*,Δ*Lk*) are time-independent, corresponding to non-equilibrium steady states (NESS), we fully characterize pathways of topology simplification mediated by type-II enzymes as network graphs having steady-state probabilities *P*\*(*K*, Δ*Lk*) and probability currents *I*[(*K*,Δ*Lk*)→(*K*’,Δ*Lk*’)] (Figures 5, 6). Our approach thus comprehensively and simultaneously addresses the kinetics of superhelix relaxation and knot resolution. One novel feature of our model is that we consider the biologically relevant case that complex knots are generated extraneously (Figure 6). Our analysis complements the work of Shimokawa and colleagues, who considered stepwise unlinking of DNA-replication catenanes by the Xer site-specific recombinase (25). Indeed, our approach can be generalized to quantitatively analyze rates of linking/unlinking during site-specific recombination and other processes.

As a starting point for our non-equilibrium model, we first investigated the equilibrium probability distribution *P*(*K*, Δ*Lk*) and free-energy landscape *F*(*K*, *Δ*Lk*) = −k_B_T* ln*P*(*K*, Δ*Lk*) to obtain the most likely relaxation path of a given DNA knot by a hypothetical topoisomerase that lacks any bias towards topology simplification and is driven only by the topological free-energy gradient. In particular, we clarify two apparently contradictory results in the literature concerning how supercoiling and knotting affect the thermodynamically most-stable topology of a circular DNA molecule. A previous study used Monte Carlo simulations to address the dependence of the topological free energy of knotted circular DNA on supercoiling and showed that non-trivially knotted species were free-energy minima for even modest, fixed values of |Δ*Lk*| (18). Moreover, complexity of the knots corresponding to the free-energy minimum increases with increasing |Δ*Lk*| (Supplementary Figure S2); thus, supercoiling favors more complex knots according to this view (18). In a study published nine years later a different team argued that in type-II enzyme action an effective linking number difference, Δ*Lke* = Δ*Lk* - 〈*Wr*〉 (*K*,nicked),is fixed instead of Δ*Lk* (cf. Figure 3), and concluded that the unknot is a universal free energy minimum, consistent with a picture in which supercoiling inhibits DNA knotting (19). The apparent contradiction is resolved by considering the full free-energy landscape of knotted supercoiled DNA, in which distributions for fixed Δ*Lk* or Δ*Lke* correspond to different paths along the contours of this landscape (Figure 4). Thus, both statements are correct depending on the particular context of the analysis.

Our non-equilibrium model recapitulates the experimental observation that type-II topoisomerases remove crossings in trefoil knots in DNA below the level expected at thermal equilibrium (8) (Figure 5). As found previously, the efficiency of unknotting strongly depends on the presence or absence of a topoisomerase-induced bend in the gate (G) segment (9, 15): a hairpin-like G segment having an induced bend of 120° gave more efficient unknotting than an unbent G segment, resulting in an 8-fold reduction of trefoil knots in the hairpin G segment case compared to a straight G segment. In addition, for our Δ*Lk*-resolved model we show that the efficiency of unknotting (the reduction factor for trefoil knots) depends strongly on the value of Δ*Lk* (Figure 5). We find that the Δ*Lk* distribution in the unknot is narrower, i.e., the DNA is less supercoiled on average in the presence of type-II enzyme activity compared to the product Δ*Lk* distribution for a type-I enzyme, in agreement with experimental results (8) (Supplementary Figure S3). The latter does not consume the energy of ATP hydrolysis and therefore generates the Δ*Lk* distribution expected at equilibrium.

Introducing an extraneous biological process that continuously converts unknotted DNAs with Δ*Lk* = 0 to a complex topological form (*K_S_*, Δ*Lk_S_*) (chosen to be (10.139^−^,−12~ in our study) at a constant rate *k_S_* leads to the following main results for the pathways of topology simplification mediated by type-II enzymes (Figure 6):

1. Only a small number of intermediate topological states contribute to the pathways, namely those that dominate the equilibrium distribution *P*(*K*|Δ*Lk*) (Supplementary Figure S2);
2. The pathways closely follow the path Δ*Lke* = Δ*Lk* − 〈*Wr*〉 (*K*, nicked) = 0 (pathways shown on the left in Figures 6A, 6B) corresponding to the minimum in the free-energy landscape *F*(*K*, Δ*Lk*) = −*k_B_T* ln *P*(*K*, Δ*Lk*) (white line on the left in Figure 4);
3. The unknotting efficiency strongly depends on the geometry of the G segment and on the degree of DNA supercoiling, being largest for a hairpin-like G segment activity in DNA for which a finite degree of supercoiling is maintained (pathway shown on the right in Figure 6A). These results suggest that only the combined effects of type-II topoisomerase activity, driving the system away from equilibrium, and increased DNA supercoiling can generate the degree of topology simplification observed in experimental measurements;
4. The dominating pathways for hairpin and straight G segments are closely similar. This surprising result can be explained by the fact that the unknotting capability of a type-II enzyme with a hairpin G segment compared to a straight G segment is enhanced mainly due to an enhanced juxtaposition probability in complex knots and enhanced probability for an unknot to remain unknotted after strand passage, as opposed to a different selection of strand passages in knotted DNA (Figure 7). In this sense, the requirement for a bent G segment acts as a topological filter. Type-II enzymes that require a hairpin G segment are not “smarter” than type-II enzymes that employ a straight G segment (the latter closely corresponding to the equilibrium situation), but rather are more active.

Other models apart from the hairpin-like G segment have considered the ramifications of “hooked” juxtapositions on topology simplification (12). The main difference between the hooked-juxtaposition model from the hairpin-like G segment model is that the enzymes bind two juxtaposed DNA segments simultaneously rather than successively. Thus the principle of both models is essentially the same, apart from the fact that hooked juxtapositions occur much more rarely than juxtapositions with a hairpin-like G segment (9, 15). Moreover, it is difficult to imagine how the enzyme could impose a geometric requirement for hooked juxtapositions on the transiently passed T segment. For this to be the case the enzyme would need to have preferential affinity for a pre-bent incoming T segment, implying also that there should be a preferred geometric orientation of this segment. We are not aware of any experimental evidence to support the latter requirement.

Results obtained in this study are based on the assumption that the affinity of type-II enzymes to bind to DNA and generate an appropriate G segment geometry is independent of the topological state (*K*, Δ*Lk*) of the DNA, in particular, independent of the degree of supercoiling. This implies that the constant *k*_0_ in Equation (2), describing the affinity and concentration of the enzyme, is assumed to be independent of the topological state (*K*, Δ*Lk*) of the DNA. Thus the constant *k*_0_ drops out in the ratio in Equation (4) so that our results are universal in the sense that they do not depend on the value of *k*_0_. However, recent experimental results suggest that type-II enzymes have a propensity to bind to DNA and form G segments in highly supercoiled DNA, presumably because the latter is strongly bent on average, thereby facilitating the formation of bent G segments (47). This effect can be implemented in our model by making *k*_0_ in Equation (2) a function of the degree of supercoiling Δ*Lke* = Δ*Lk* − 〈*Wr*〉 (*K*, nicked).

It has long been argued that for thermodynamic reasons type-II enzyme action requires the energy of ATP hydrolysis to move the system out of topological equilibrium. Bates *et al*. have argued that only a small portion of the free energy gained from ATP hydrolysis is needed to achieve topology simplification (48). In a study of *E. coli* topoisomerase-IV mutants, Lee *et al*. found that the extent of topoisomerase II-induced DNA bending in the substrate DNA G segment, but not DNA binding, was correlated with ATP-hydrolysis activity (49). Thus, a relevant question in this context is at which point during the strand-passage reaction the energy gained from ATP hydrolysis is used by the enzyme, and for what purpose. Even without ATP hydrolysis the enzyme can bind to DNA and perform strand passage (50, 51); this implies that these steps are essentially driven by the free-energy gradient so that the enzyme-DNA complex after strand passage should be very stable. Thus it has been proposed that ATP hydrolysis serves to release the energy necessary for dissociating the stable enzyme-DNA complex after strand passage, and resetting the original conformation of the protein (14, 52) (Figure 1A). Other studies suggested that two ATP molecules are hydrolyzed sequentially before and after strand passage, respectively (53, 54). It would be interesting to address these questions by modeling the enzymatic reaction in terms of graphs on networks formed by chemical and conformational states of the enzyme-DNA complex, similar as has been recently done for molecular motors and other nanomachines (55–58).

## ACKNOWLEDGEMENT

We thank Keir Neuman and Cristian Micheletti for communicating results to us in advance and for helpful discussions.

## Supplementary Material

### S1. JOINT PROBABILITY DISTRIBUTION, *P*(*K*, Δ*Lk*)

The free energy landscape *F*(*K*, Δ*Lk*) = −*k_B_T* ln *P*(*K*, Δ*Lk*) shown in Figure 4 was obtained by calculating the joint probability distribution *P*(*K*, Δ*Lk*) of knot type *K* and linking number difference Δ*Lk* for 6-kbp DNA using the DNA model and Monte Carlo (MC) simulation procedure described in the main text. To calculate *P*(*K*,Δ*Lk*) we used the relation [1]

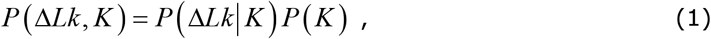

where *P*(Δ*Lk*|*K*) is the conditional distribution of Δ*Lk* for given *K* and *P*(*K*) is the distribution of *K* for torsionally unconstrained (nicked) DNA.

*P*(*K*) in Equation (1) was determined by MC simulations from the frequency of occurrence of knot types *K* in equilibrium segment-passage (ESP) ensembles containing 10^6^ saved conformations, where the simulation period between saved conformations was 1000 MC moves. Since the probability of occurrence of any particular knot decreases exponentially with its complexity [2] we used the method of restricted ESP ensembles [3] to accurately determine *P*(*K*) for complex knots (Table S1, Figure S1). This method uses restricted ensembles in which one or more dominating knot types are excluded so that less dominant knot types occur with higher frequency. Using the relation *P*(*B*)/*P*(*A*) = *P*’(*B*)/*P*’(*A*) for the probabilities of occurrence of knot types *A*, *B* in the unrestricted ensemble, ESP, and restricted ensemble, ESP′, respectively, one obtains the probability *P*(*B*) of a knot *B* that occurs with low frequency in ESP but with sufficiently high frequency in ESP’ as

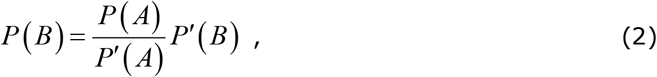

where the knot *A* serves as link between ensembles ESP and ESP′. If the probability of a knot *C* is too low even in the restricted ensemble ESP’ one may iterate the procedure by including an even more restricted ensemble ESP″, resulting in

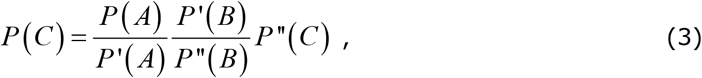

where knot *A* serves as link between ESP and ESP′, and knot *B* as link between ESP′ and ESP″ (Figure S1). In our calculations we considered ensembles in which no knot types were excluded (ESP) as well as the restricted ensembles ESP′ = ESP−{0.1} and ESP″ = ESP′−{3.1, 4.1, 5.1, 5.2} (where for the chiral knots 3.1, 5.1, 5.2 both the left and right-handed forms were excluded). The knots *A* = 3.1^−^ and *B* = 6.1^−^ served as links between ensembles according to Equations (2), (3). *P*(*K*) for *K* = 0.1, 3.1^−^, 4.1 was obtained from ESP, for 5.1^−^, 5.2^−^ from ESP′ using Equation (2), and for the remaining knots shown in Table S1 from ESP″ using Equation (3).

*P*(Δ*Lk*|*K*) in Equation (1) was calculated using the relation [1]

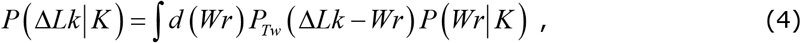

where *P*(*Wr*|*K*) is the distribution of writhe *Wr* for given knot type *K* and *P_Tw_* (Δ*Tw*) is the distribution of twist Δ*Tw* in torsionally relaxed (nicked) DNA. *P_Tw_* (Δ*Tw*) is assumed to be of Gaussian form 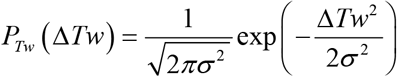, with variance *σ*^2^ = *N*/(4*π*^2^*cM_tw_*), where here *N* = 200 and *c_tw_* = 7.243, and White’s equation in the form Δ*Tw* = Δ*Lk* − *Wr* was used (see main text). The distribution *P*(*Wr*|*K*) in Equation (4) was obtained by MC simulations of torsionally unconstrained chains with fixed knot type *K* . The resulting mean values 〈*W*〉(*K*) for various knot types used in the main text are shown in Table S1.

**Figure S1.**
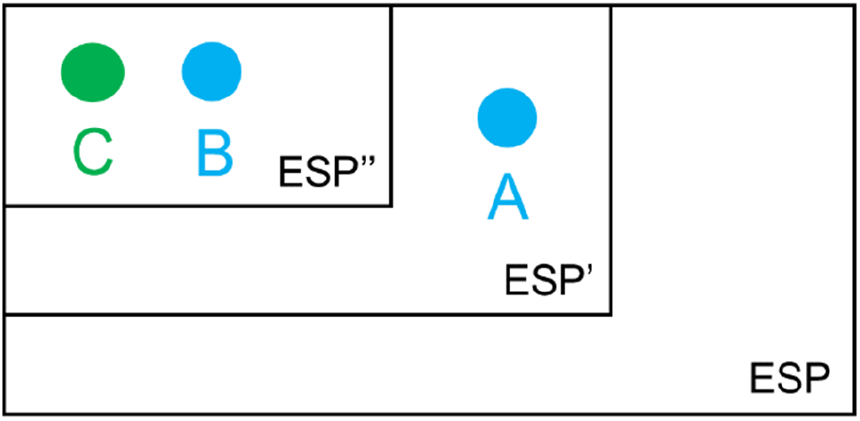
Schematic illustration of the method of restricted ensembles to obtain the probability *P*(*C*) of a complex knot *C* (green dot) which occurs with very low frequency in the unrestricted ensemble ESP but with sufficient frequency in the restricted ensemble ESP″. Knots *A* and *B* (blue dots) link ESP with an intermediate restricted ensemble ESP′ and ESP′ with ESP″, respectively (see Equation (3)).

**Table S1.**
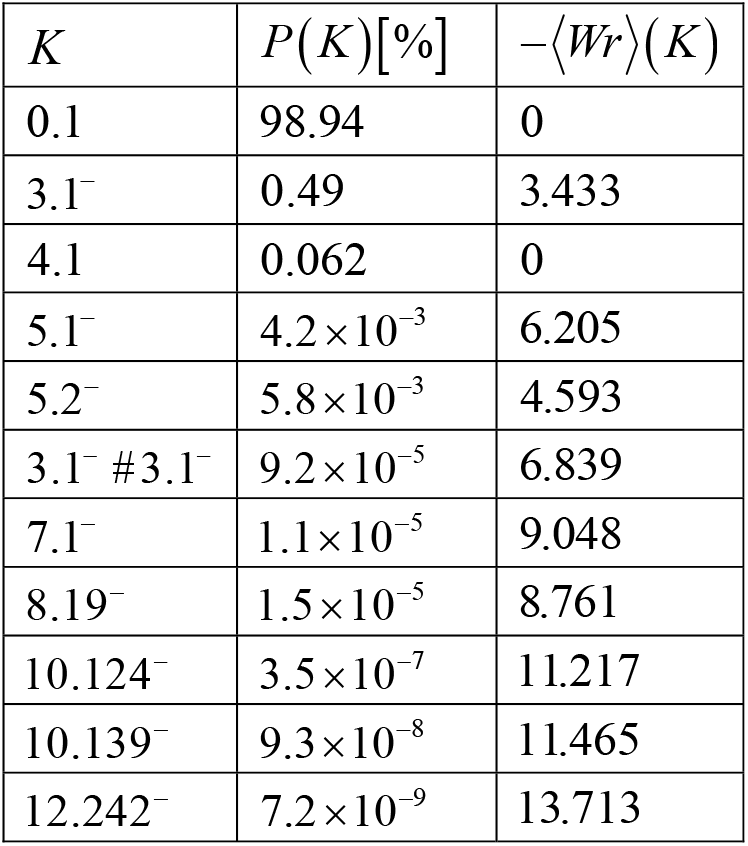
Equilibrium probabilities *P*(*K*) in per cent and mean writhe 〈*Wr*〉(*K*) in ESP ensembles of various torsionally unconstrained knots *K* for 6-kbp DNA’s (*N* = 200 segments).

| $K$ | $P(K)[\%]$ | $-\langle Wr \rangle(K)$ |
| --- | --- | --- |
| 0.1 | 98.94 | 0 |
| 3.1<sup>-</sup> | 0.49 | 3.433 |
| 4.1 | 0.062 | 0 |
| 5.1<sup>-</sup> | $4.2 \times 10^{-3}$ | 6.205 |
| 5.2<sup>-</sup> | $5.8 \times 10^{-3}$ | 4.593 |
| 3.1<sup>-</sup> # 3.1<sup>-</sup> | $9.2 \times 10^{-5}$ | 6.839 |
| 7.1<sup>-</sup> | $1.1 \times 10^{-5}$ | 9.048 |
| 8.19<sup>-</sup> | $1.5 \times 10^{-5}$ | 8.761 |
| 10.124<sup>-</sup> | $3.5 \times 10^{-7}$ | 11.217 |
| 10.139<sup>-</sup> | $9.3 \times 10^{-8}$ | 11.465 |
| 12.242<sup>-</sup> | $7.2 \times 10^{-9}$ | 13.713 |

### S2. CONDITIONAL PROBABILITY DISTRIBUTION, *P*(*K*, Δ*Lk*)

**Figure S2.**
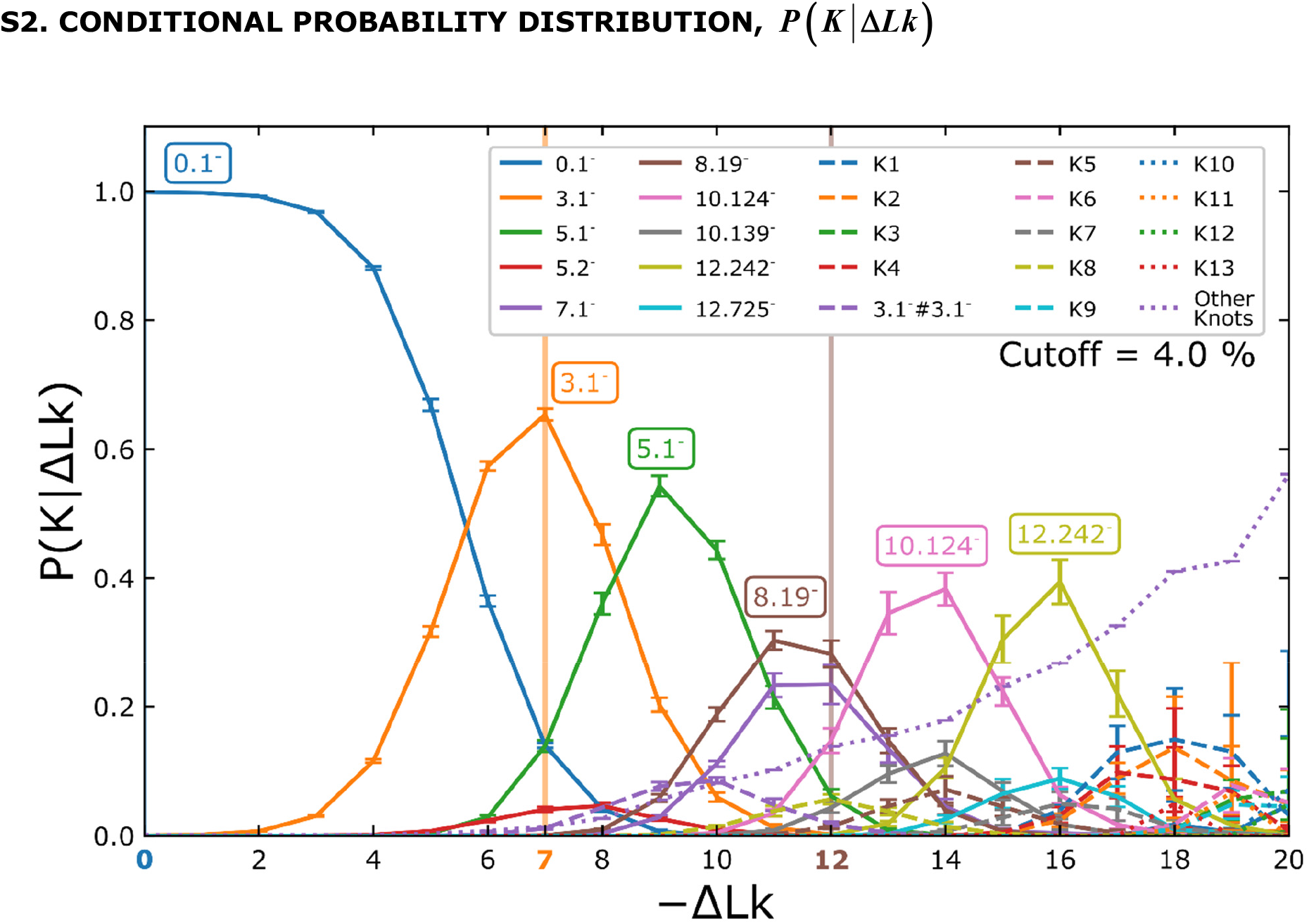
Conditional probability distribution *P*(*k*|Δ*Lk*) of knot types *K* with given linking number difference Δ*Lk* for 6-kbp DNAs. *P*(*K*| Δ*Lk*) was calculated by MC simulation of ESP ensembles with fixed linking number Δ*Lk* and measuring the frequency of occurrence of knot types *K* . Only curves with *P*(*k*| Δ*Lk*) > 0.04 are shown. For any fixed, small value of Δ*Lk* only a few knot types *K* dominate the distribution. Conversely, for −Δ*Lk* > 18 the distribution degenerates and many different knot types *K* contribute to *P*(*K*|Δ*Lk*). The values −Δ*Lk* = 7 and −Δ*Lk* = 12 indicated by orange and brown vertical lines, respectively, refer to Figure 4 in the main text. Knots K1-K13 have more crossings than knots in any tables available to us.

### S3. NARROWING OF THE STEADY-STATE DISTRIBUTION, *P*^*^(Δ*Lk*|0.1)

**Figure S3.**
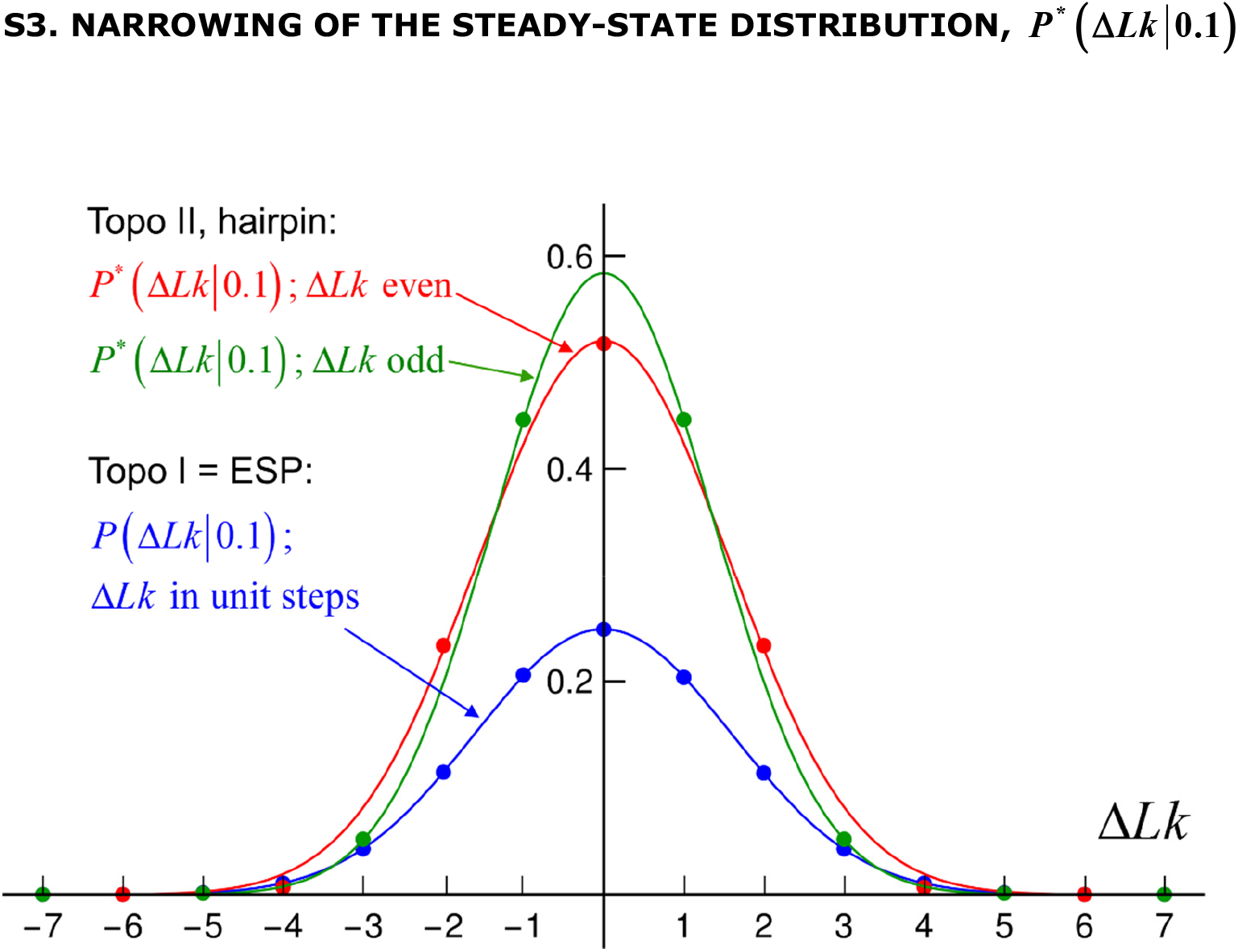
Steady-state distribution *P*\*(Δ*Lk*|0.1) for the unknot 0.1 in the presence of type-II enzymes modeled as hairpin-like G-segment. Because type-II enzymes change Δ*Lk* in steps of 2, the distributions *P*\*(Δ*Lk*|0.1) for even and odd values of Δ*Lk* are disjunct (red and green dots, respectively). Also shown is the distribution *P*(Δ*Lk*|0.1) for type-I enzymes, which essentially corresponds to an equilibrium (ESP) distribution, changing Δ*Lk* in steps of 1. The solid lines are least-square Gaussian fits to the discrete distributions. The distributions *P*\*(Δ*Lk*|0.1) and *P*(Δ*Lk*|0.1) are used to calculate the topology simplification factor *TSF* quantifying the narrowing of the Δ*Lk* - distribution in the presence of type-II enzymes and ATP relative to type-I enzymes.

Variance of Δ*Lk* for type-II enzyme with hairpin G-segment for even and odd values of Δ*Lk* :

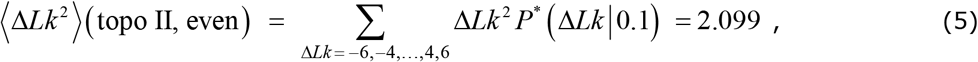

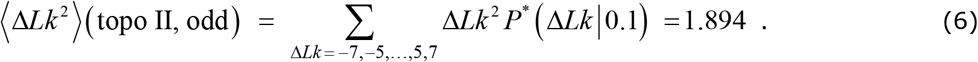

Variance of Δ*Lk* for an equilibrium (ESP) distribution with Δ*Lk* in unit steps:

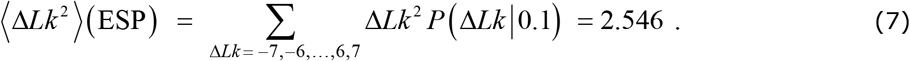

Resulting topology simplification factor *TSF* = s.d.(Δ*Lk*,topo II)/s.d.(Δ*Lk*,ESP) for the standard deviation 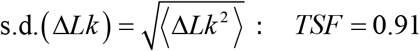 for Δ*Lk* even and *TSF* = 0.86 for Δ*Lk* odd.

### S4. DEPENDENCE OF *P*(*a*) AND *I_ab_* ON SOURCE RATE *k_S_*

**Figure S4.**
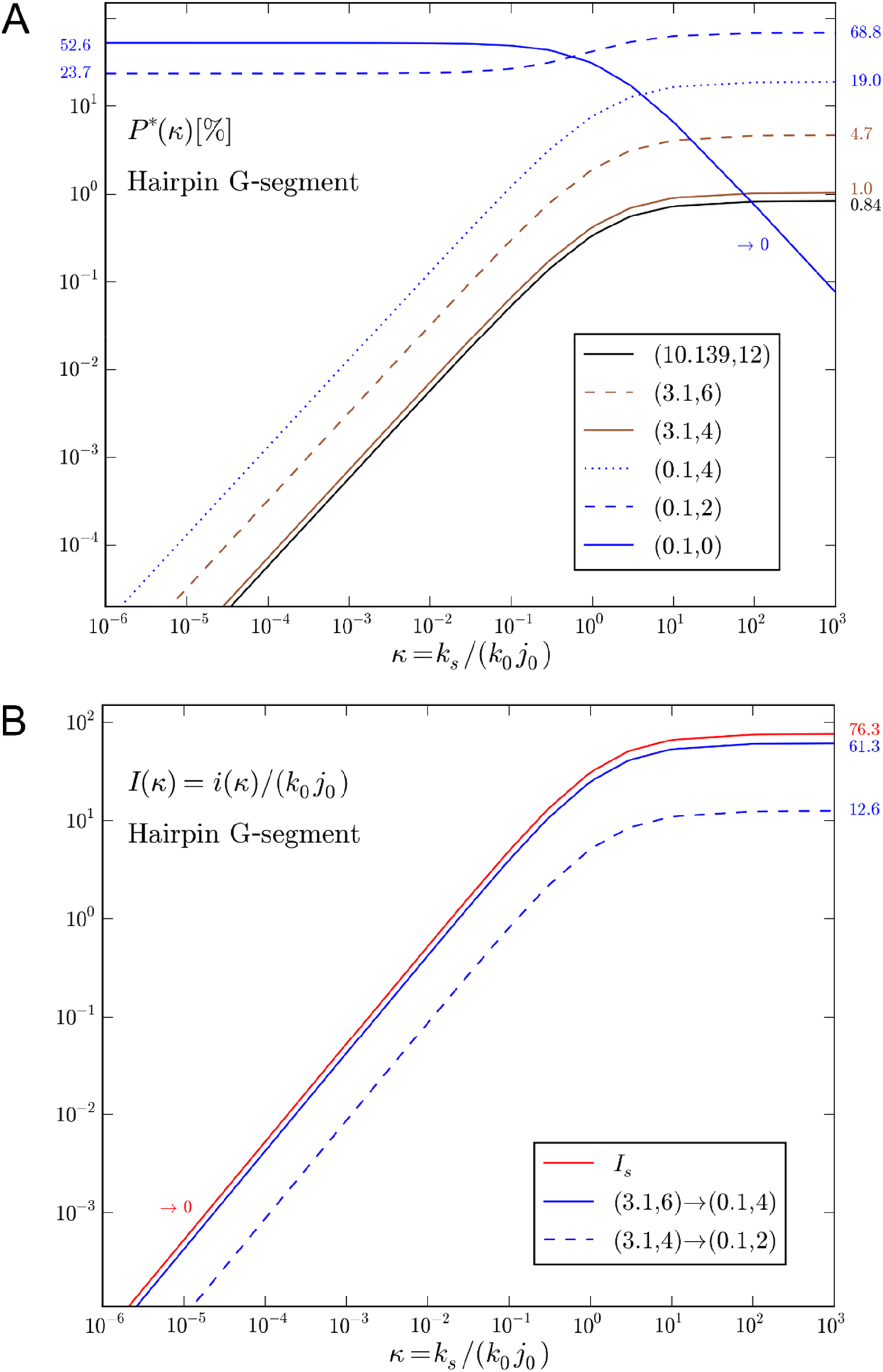
(**A**) Steady-state probabilities *P*\*(*a*) in per cent and (**B**) normalized probability currents *I_ab_* as function of the parameter *ĸ* = *k_S_*/(*k*_0_*j*_0_) for type-II enzyme with hairpin G-segment for various topological states *a* = (*K*, Δ*Lk*), *b* = (*K*’, Δ*Lk*’) (cf. Computational Methods and Figure 6A). The curves show the crossover behavior of *P*\*(*a*) and *I_ab_* between the limits *k_S_* = 0 (corresponding to the absence of an external process that generates a source state *a_S_*) and *k_S_* → ∞. Corresponding limit values of *P*\*(*a*) and *I_ab_* are indicated by numbers on the left and right sides of the figures, respectively. The steady-state probability of the originating state (0.1,0) in (**A**) vanishes as *P*\*(0.1,0) ~ 1/*k_S_* for large source rate *k_S_* since this state is depleted by the process that generates DNAs in the source state *a_S_* = (10.139^−^,−12). For all other states *a* the steady-state probabilities approach finite values 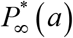 in the limit of a large source rate *k_S_* . Likewise, all probability currents *I_ab_* approach finite values 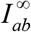 in the limit of large source rate *k_S_*, including the source current *I_S_*.

**Figure S5.**
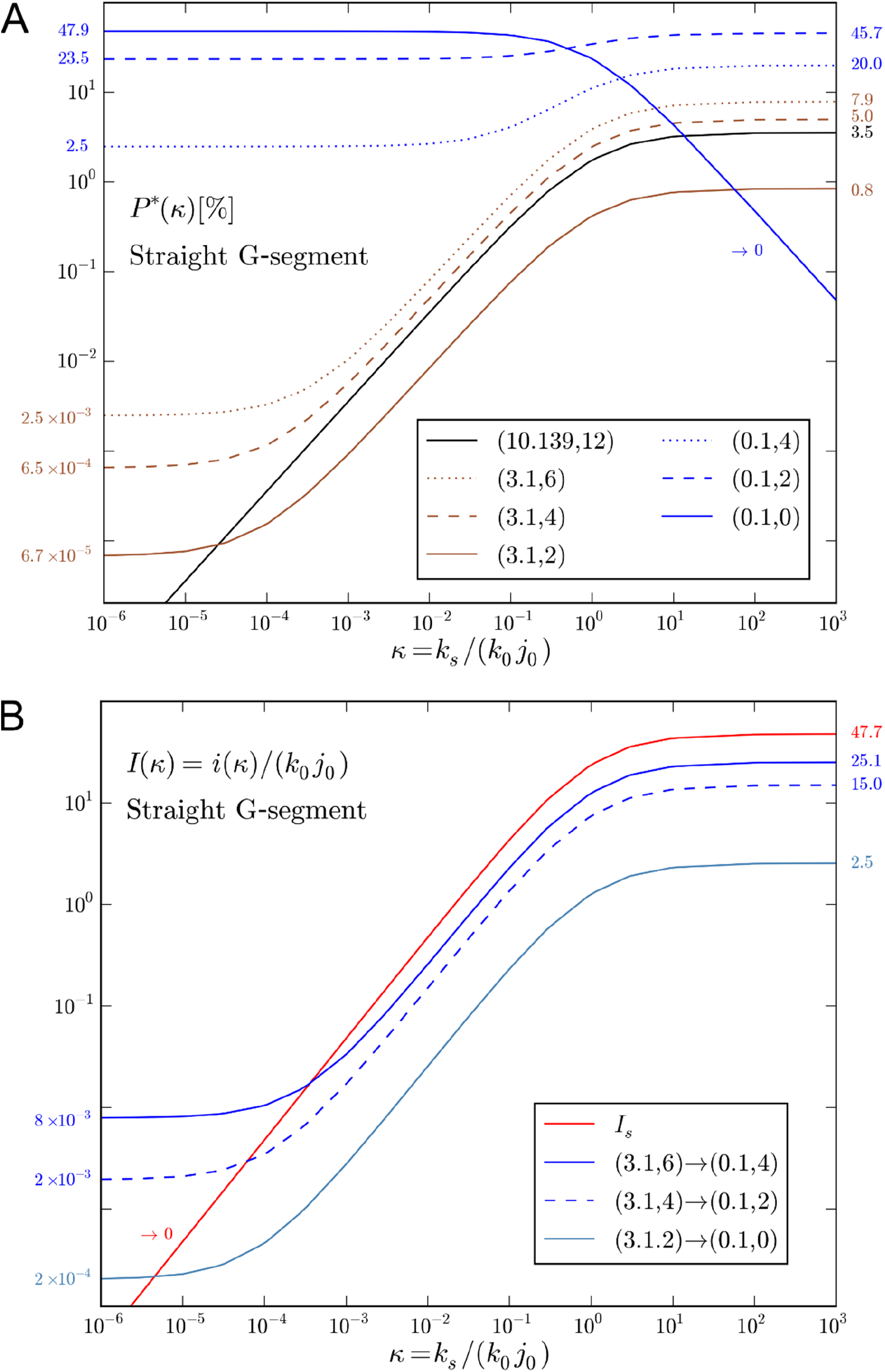
Same as Figure S4 for type-II enzyme with straight G-segment (cf. Figure 6B).

### S5. UNIVERSALITY OF STEADY-STATE PROBABILITIES, *P*\*(*K*)

**Figure S6.**
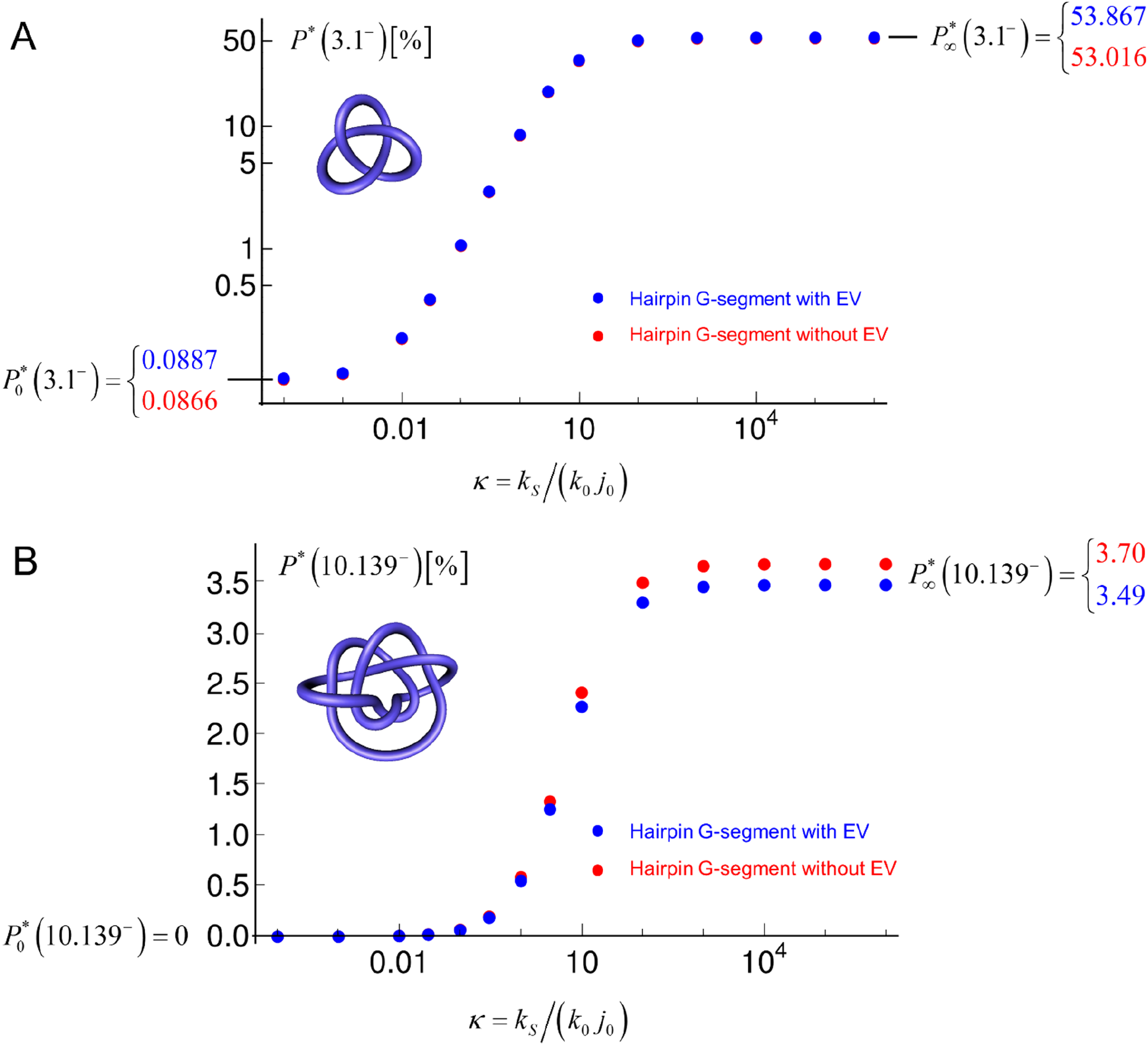
Steady-state probabilities *P*\*(*K*) in per cent for (**A**) *K* = 3.1^−^ and (**B**) *K* = 10.139^−^ for torsionally unconstrained (nicked) 6-kbp DNAs as function of the parameter *ĸ* = *k_S_*/(*k*_0_ *j*_0_) for type-II enzyme with hairpin G-segment. Shown are results for two cases: (i) the four segments forming the hairpin G-segment (shown red in Figure 2A) interact with the segments of the rest of the chain (including a potential T-segment) with the same excluded volume (EV) interaction as the segments of the rest of the chain (blue dots); (ii) the four segments forming the hairpin G-segment do not interact with the rest of the chain by an EV interaction (red dots). Thus, the G-segment in case (ii) does not repel the segments of the rest of the chain by an EV interaction, and in this sense is more active than the G-segment in case (i). As a result, the juxtaposition probability *j*_0_ for the unknot in our simulations is about 10 times larger in case (ii) than in case (i): *j*_0_ (without EV) = 0.00156, *j*_0_(with EV) = 0.000162 . Nevertheless, the curves *P*\* (*K*) as functions of the parameter *ĸ* = *k_S_ļ* (*k*_0_ *j*_0_) collapse on single curves in both cases within numerical error, which shows that *P*\* (*K*) is independent of whether the G-segment is modeled with or without EV interaction. More generally, *P*\*(*K*) is expected to be independent of any property of the enzyme that determines its overall activity regardless of the topological state of the DNA. In this sense, *P*\*(*K*) as function of *κ* is universal. A key requisite for this universality is that the enzyme is much smaller than the DNA, so that the enzyme itself cannot probe the topological state of the DNA. In (**A**) we use a log-log scale to show that *P*\*(3.1^−^) approaches a finite value 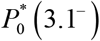 in the limit *k_S_* → 0 (corresponding to the absence of an external process that generates the knot 3.1^−^) as indicated on the left side of the figure. In (**B**) we use a linear scale for the *y*-axis because *P*\*(10.139^−^) → 0 for *k_S_* → 0 . The finite limits 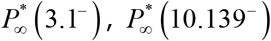 for *k_S_* → ∞ are indicated on the right sides of figures (**A**), (**B**). We attribute the small deviation between the curves for cases (i) and (ii) by the fact that the enzyme in our simulation, albeit being small, has a finite size compared with the rest of the chain, which has a larger effect on the knot 10.139^−^ than on 3.1^−^ since the former is more compact on average than the latter.

